# Identification of Conserved Evolutionary Trajectories in Tumors

**DOI:** 10.1101/2020.03.09.967257

**Authors:** Ermin Hodzic, Raunak Shrestha, Salem Malikic, Colin C. Collins, Kevin Litchfield, Samra Turajlic, S. Cenk Sahinalp

## Abstract

**Motivation:** As multi-region, time-series, and single cell sequencing data become more widely available, it is becoming clear that certain tumors share evolutionary characteristics with others. In the last few years, several computational methods have been developed with the goal of inferring the subclonal composition and evolutionary history of tumors from tumor biopsy sequencing data. However, the phylogenetic trees that they report differ significantly between tumors (even those with similar characteristics).

**Results:** In this paper, we present a novel combinatorial optimization method, CONETT, for detection of recurrent tumor evolution trajectories. Our method constructs a consensus tree of conserved evolutionary trajectories based on the information about temporal order of alteration events in a set of tumors. We apply our method to previously published datasets of 100 clear-cell renal cell carcinoma and 99 non-small-cell lung cancer patients and identify both conserved trajectories that were reported in the original studies, as well as new trajectories.

**Availability:** CONETT is implemented in C++ and available at https://github.com/ehodzic/CONETT.

## 1 Introduction

Cancer is a disease initiated by somatic genomic alterations. Accumulation of such alterations drives the progression of a cancer to an advanced form. As the tumor grows, new subpopulations of the tumor cells emerge with distinct genomic alteration profiles. After subsequent rounds of selection and expansion, these subpopulations give rise to substantial intra-tumor heterogeneity, which is arguably one of the main challenges in cancer management and treatment.

As multi-region, time-series and single cell sequencing data become more widely available, it is becoming clear that certain tumors share evolutionary characteristics with others. With new computational methods to identify recurrent cancer progression patterns from *multi-dimensional* tumor sequencing data, it may become possible to predict the likely course of evolution and perhaps an effective treatment strategy for certain cancer types [15, 23, 8]. E.g. novel approaches for targeted immuotherapy [6] could be developed into combinatorial strategies that simultaneously target multiple subclonal populations if a comprehensive characterization of recurrent subclonal expansion patterns could be established. Unfortunately, elevated levels of intra-tumor heterogeneity, with intricate subclonal mutational landscapes dominated by inconsequential passenger alterations, may make it difficult to discriminate predictable tumor progression *signals* from *noise* introduced by these statistically insignificant subpopulations.

In the last few years, several computational methods have been developed with the goal of inferring the subclonal composition and/or evolutionary history of tumors from tumor biopsy sequencing data including [10, 7, 29, 24, 11, 17, 30, 35, 9, 12, 25, 26, 36]. Phylogenetic trees for representing tumor evolution processes reported by such methods provide (at least one of) the likely relative temporal order of genomic alteration events. Unfortunately, the trees they report for many tumor samples differ significantly from other tumors with similar characteristics. In contrast, some recent studies have been able to recapitulate tumor evolution more accurately through long-term clinical followup [34, 1, 31]. These studies were able to find evidence of recurrent progression patterns involving two or more co-occurring genomic alterations. Since, until recently, no computational tool was purposefully developed to identify recurrent patterns of tumor evolution across multiple tumors, these studies typically relied on manual inspection of all possible sequences of alteration events, starting from the root of the tumor phylogeny (representing germline), all the way to leaf nodes [31]. Recently, methods have been developed that cluster tumor phylogenetic trees into groups of closely related trees with respect to edge similarity/distance measures [27, 5, 2]. In particular, the REVOLVER method [5] employs a maximum-likelihood learning strategy to construct a joint hidden tree model for all tumors in a cohort, which is then used to infer an individual tree depicting the temporal order of clonal alterations in each individual tumor (this particular aspect of the method is later improved upon by [20]). This is followed by a hierarchical clustering of the trees for the purpose of detecting edges shared within each cluster. We note here that REVOLVER neither constructs likely evolutionary *trajectories* (which may be done manually to a degree [5] via the use of shared edges) nor does it look into whether the trajectories are conserved in a significant portion of tumors.

In parallel to the effort summarized above, the problem of finding common sets of genomic alterations in cross-sectional tumor data has been investigated by several computational approaches. These approaches typically employ molecular interaction networks to identify *connected subnetworks* of genes that have been subjected to somatic alterations, that are *shared* across tumor samples in the sense that at least a fixed number (e.g. one) of the genes in the subnetwork is altered in each tumor sample [33, 22, 21, 32, 3, 4, 16]. Recently, cd-CAP [14] took a stricter approach, requiring preservation of every alteration in the subnetwork in at least a user defined number of tumor samples in a cohort, introducing the notion of a *conserved alteration pattern*. Through cd-CAP it has become possible to detect sizable networks of genes, altered in a similar way across tumor samples. However none of these approaches consider the *temporal order of alterations* to make inference about evolutionary processes common in certain tumor types.

In this paper, we present a novel computational method, CONETT (CONserved Evolutionary Trajectories in Tumors), for combinatorial detection of recurrent tumor evolution trajectories. CONETT considers as input a partial temporal ordering of alteration events for each tumor in a given set of tumors; for any tumor the input is given in the form of a directed graph where each alteration event is represented by a distinct node. Given a pair of alteration events *a* and *b*, one of the following must be true for each tumor: (i) a directed edge from *a* to *b* only (alternatively from *b* to *a* only) indicates that *a* is an ancestor of *b*, (ii) a directed edge from *a* to *b* as well as a directed edge from *b* to *a* (or a bidirectional *a*-*b* edge), indicates that the two events are known to have an ancestor-descendant relationship but their specific ordering is unknown, (iii) an “anti-edge” between *a* and *b* indicates that the two events belong to two distinct lineages, or (iv) a “don’t care” edge between *a* and *b* indicates that either *a* or *b* or both are not observed in that tumor or no information is available with respect to their ordering.^1^ Given such input, and a “root” (e.g. driver) alteration event *s*, CONETT constructs a “consensus” phylogeny tree on a large subset of tumors whose topology “captures” the ancestor-descendant relationship for the largest possible number of event pairs in these subset of tumors. A path from the root of that phylogeny to any other event is said to form an “evolutionary trajectory” of alteration events, “conserved” in this subset of tumors.

In its most stringent setup, CONETT constructs a consensus tree (rooted at the germline “event” *g*) of all tumors in the cohort with maximum total node “depth” (distance from the root), under the constraint that if event *a* is an ancestor of event *b* in the consensus tree, then in *each* individual tumor in which *b* has been observed, there must exist a directed edge from *a* to *b*. As a result, not all alteration events can be a part of the consensus tree. Thus CONETT inferred consensus tree maximizes the number of ancestor-descendant orderings between pairs of alteration events that do not conflict with those orderings observed in any individual tumor graph. In a more general setup, CONETT can relax this constraint for a small fraction of tumor graphs. It also allows the user to specify a node *s* ≠ *g* as the “driver” event.

We have applied CONETT to the TRACERx clear-cell renal cell carcinoma (ccRCC) data set [31] involving 100 tumors, as well as TRACERx non-small-cell lung cancer (NSCLC) data set [18] involving 99 tumors, and identified a number of significantly conserved evolutionary trajectories which involve sequence-altered genes and copy number alteration events – not reported in the original studies.

## 2 Methods

Let 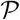 be a set of tumors for which it is possible to infer (possibly through the use of time series, multi-region, single cell or single molecule sequencing data) a partial ancestral ordering of alteration events, labelled by a gene or chromosome and the type of alteration affecting it (e.g. somatically altered single gene or copy number altered entire chromosomal arm) ^2^. Let 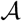 be the set of all alteration events that are observed in at least one tumor 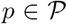. For a given tumor 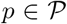, we define a *tumor graph G_p_* as a graph that has a node for each alteration event in 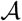 (note that *G_p_* includes a node for each event in 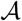, but not every node in *G_p_* corresponds to an event observed in *p*). For each pair of distinct nodes *a, b* ∈ *V* (*G_p_*) exactly one of the following must be true: the set of edges *E*(*G_p_*) contains (i) either a directed edge from *a* to *b* only or from *b* to *a* only, indicating that *a* is an ancestor of *b* (or alternatively, that *b* is an ancestor of *a*); (ii) a directed edge from *a* to *b* as well as a directed edge from *b* to *a*, indicating that the two events are known to have an ancestor-descendant relationship but the specific ordering is unknown; (iii) an “anti-edge” between *a* and *b*, indicating that the two events belong to two distinct lineages; or (iv) a “don’t care” edge between *a* and *b*, indicating that alteration events corresponding to either one or both of the nodes are not observed in that tumor or no information is available with respect to their ordering. We require that each tumor graph should be transitive with respect to its directed edges, i.e. if there are directed edges from *a* to *b* and from *b* to *c* then there must exist a directed edge from *a* to *c*. Thus, edges of a tumor graph *G_p_* capture the complete available information about partial ancestor-descendant ordering of alteration events in tumor *p*.

We say that an ordered set of nodes *e* = (*v*_1_,…,*v_k_*) represents an *evolutionary trajectory* that is *conserved* in a tumor graph 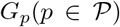 if for each pair of nodes *v_i_, v_i_*_+1_ in *e* there is a directed edge from *v_i_* to *v_i_*_+1_. In its most stringent setting, CONETT builds a tree *T* that includes ancestor-descendant relationships observed in *all* tumors in 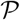 - i.e. the path from the root to any given node *v* in *T* is conserved in all tumor graphs *G_p_* for which the alteration event corresponding to *v* is observed in tumor *p*. Among such trees *T*, CONETT’s objective is to compute the tree 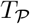 that maximizes the total path length from the root to every other node *v*; we call this tree, the “consensus” tree for 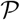, and each path in 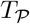 from the root to any other node a “conserved evolutionary trajectory”. Intuitively, the consensus tree maximizes the number of orderings between alteration event pairs. We note here that we represent the “germline” as a special (pseudo) alteration event *g* from which each *G_p_* has a directed edge to every other node *v*. In this stringent setting, the root of *T* is specified to be *g*, and thus *T* will include each event in 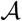 as a node - sometimes as a singleton leaf, with *g* as its parent.^3^ Singleton leaves can later be filtered out as they reveal no new information.

In a more general setting (which we use for all our experimental results), CONETT offers the ability to identify longer evolutionary trajectories that are conserved only in a subset of tumor graphs in 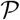, by relaxing the constraint that each trajectory should be conserved in every tumor graph. Additionally, this setting allows tumor graphs to be non-transitive (for reasons explained later in this section). Finally, this general setting allows the user to specify as the root, any alteration/node 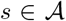, to be identified as the “driver” alteration event. For every other event *v*, let 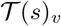, denote the set of all tumor graphs *G_p_*, where there is a directed path from *s* to *v*. We then formulate the following problem.

### Maximum Conserved Evolutionary Trajectory Tree problem (MCETT)

Construct a tree 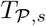 rooted at *s* with the maximum total path length from *s* to every other node *v*, such that every path (*s, u*_1_,…,*u_k_, v*) in 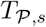 is conserved in at least 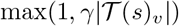 tumor graphs in 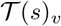, for a user defined *γ* ∈ (0, 1].

Note that in order to achieve maximum total path length (i.e. node “depth”), the solution to the *MCETT* problem necessarily includes all nodes *v* for which 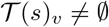. CONETT solves *MCETT* problem through an ILP formulation as described in Section 2.1. However if *s* is set to be the germline node and *γ* = 1, the solution to the *MCETT* problem is the consensus tree we defined above for the most stringent setting of CONETT and for this case we describe a simple polynomial time algorithm below.

### A Polynomial Time Algorithm for MCETT problem for *γ* = 1

Let *s* denote the root alteration event (not necessarily representing the germline). For any alteration/node 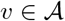, let 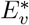 denote the set of edges (*u, v*), conserved in every tumor graph in 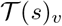 and let 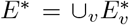. The solution to the *MCETT* problem for this setting is then the spanning tree of the nodes in *E*\* in which the sum of distances between root *s* and every other node *v* is maximum possible. It is possible to compute this spanning tree through a simple depth first search strategy as shown in Supplementary Methods.

For the more general setting of *MCETT* where *γ* < 1, the *maximum depth spanning tree* strategy above could be used as an efficient heuristic that works reasonably accurately for higher values of *γ*. However it is possible to solve *MCETT* exactly for any value of *γ* through an integer linear programming formulation as described below in Section 2.1. In this formulation, we do not differentiate (i) node pairs with a single directed edge from (ii) those with edges in both directions. We also do not differentiate (iii) node pairs with an anti-edge from (iv) those with a don’t care edge. We show how to differentially “incentivize” the first two edge types towards positive contributions, and “penalize” the next two edge types towards negative contributions within the objective of this ILP formulation later in the paper.

### 2.1 ILP formulation for solving the Maximum Conserved Evolutionary Trajectory Tree problem

Given a set of tumor graphs 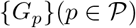, a root alteration *s* and a value of the parameter *γ* ∈ (0, 1], the goal is to construct a tree *T* as the solution to the Maximum Conserved Evolutionary Trajectory Tree problem. First, for each alteration event 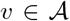, CONETT finds the maximum-size subset 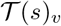 of all tumor graphs in which there is a directed path from *s* to *v* in each 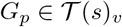. Then CONETT forms an *evolutionarily conserved alteration set* 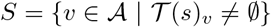, which is the set of all nodes *v* for which 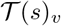 is non-empty and thus the node must be included in the solution tree.

Given an evolutionarily conserved alteration set *S*, CONETT’s ILP formulation in Figure 2 specifies the tree 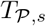 built upon the node set *S*, with *s* as its root, that represents the maximum node depth tree in which each path represents a conserved evolutionary trajectory with respect to the value of *γ*. In the ILP formulation, binary variable *X_u,v_*, defined for each pair of nodes *u, v* ∈ *S*, is set to 1 if (*u, v*) is a tree edge of 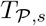, and set to 0 otherwise. For each tumor graph *G_p_* and node *v* ∈ *S*, variable 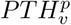 is set to 1 if the exact path in 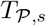 leading from *s* to *v* is conserved is tumor graph *G_p_*. Lastly, for each node *v* ∈ *S*, the variable *D_v_* represents the distance between *s* and *v* in 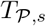. The objective of the ILP is to maximize the sum of distances from the root to all nodes in *S*.

**Figure 1:**
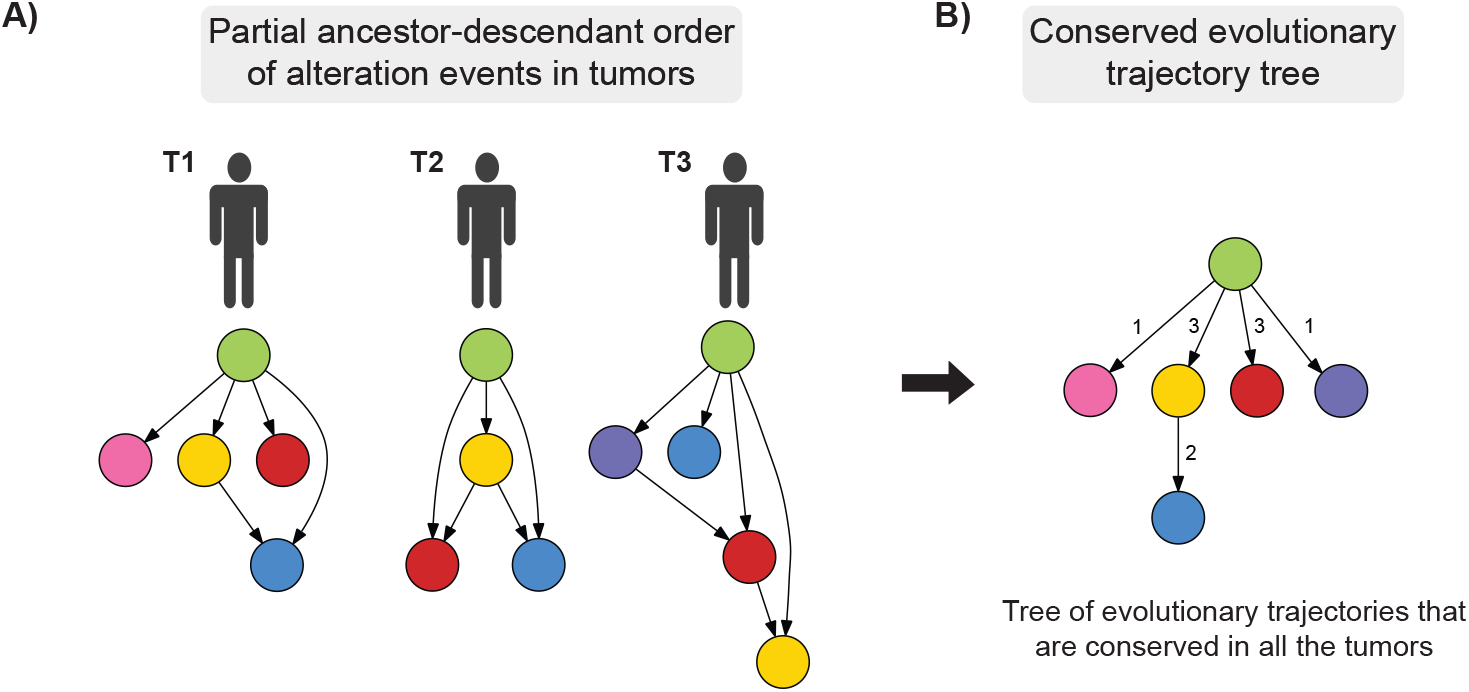
Overview of CONETT’s framework. The method takes as its input a partial ancestor-descendant ordering of alteration events across a number of tumors and produces a tree of conserved evolutionary trajectories stemming from a common root event which maximizes the number of ancestor-descendant ordered pairs of nodes. **A) Input**. Tumor graphs represent partial ancestor-descendant order of alteration events in tumors. In this particular example, any pair of events that do not have an actual edge between them in a tumor should be thought to have an “anti-edge”; similarly any event not present in a tumor should be thought to have a “don’t care” edge to all events present in that tumor. **B) The Conserved Evolutionary Trajectory Tree**. In the most stringent setting CONETT computes the maximum total depth “consensus” phylogeny tree where the ordering of alteration events do not conflict with the partial orderings of alteration events in any of the tumors.

**Figure 2:**
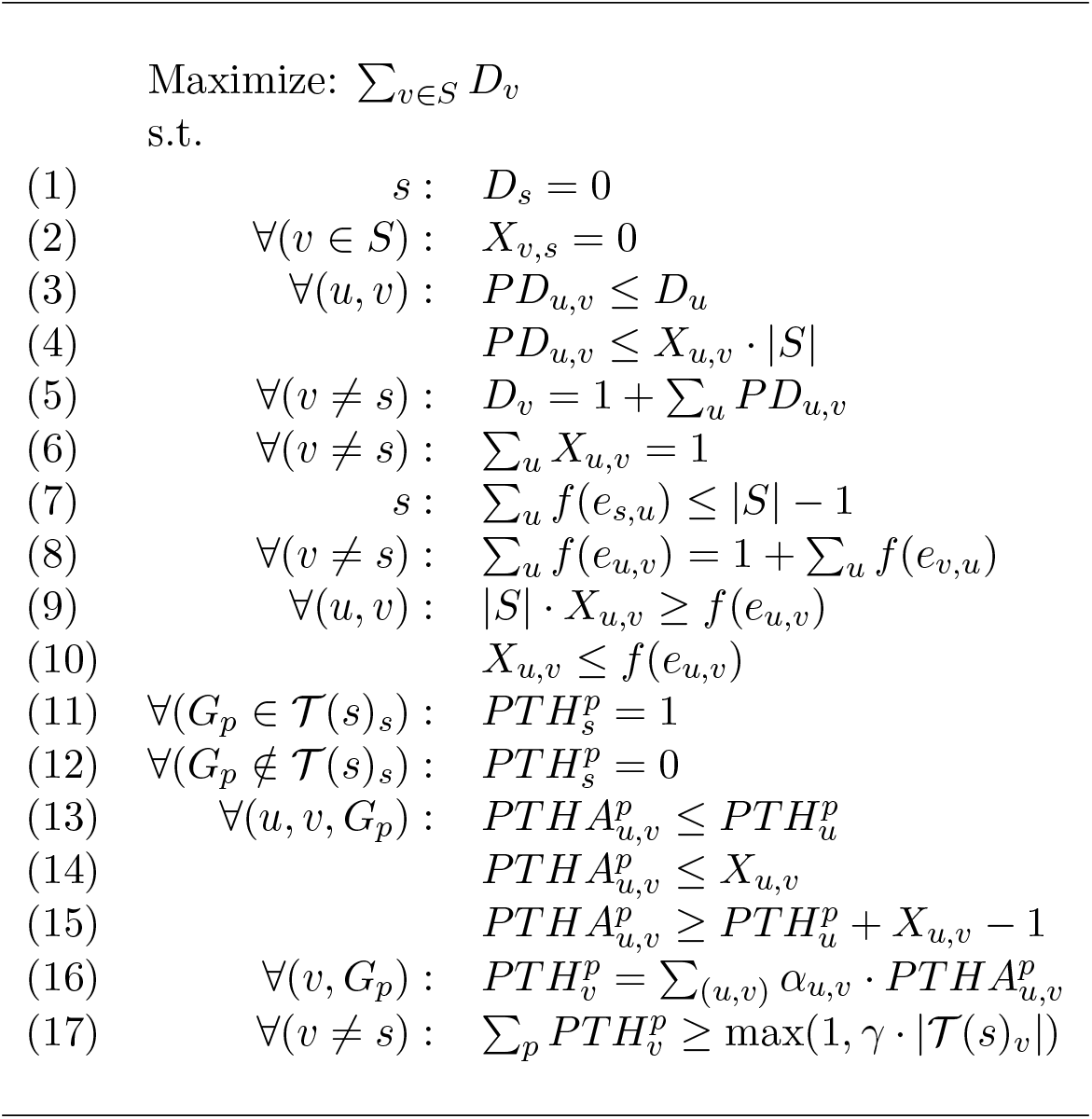
The ILP formulation for inferring the conserved evolutionary trajectory tree

Now we describe each of the constraints in the ILP formulation. First, the root node *s* is set to have distance 0 by constraint (1), and it cannot be a child of any other node due to constraint (2). Every other node *v* has distance that is equal to the distance of its parent node *u* increased by 1. The edge to node *v* from its parent *u* is determined by auxiliary variables *PD_u,v_*: Due to the constraint (4), *PD_u,v_* is set to 0 if *u* is not the parent of *v*; The maximization in the objective function, combined with the constraints (3) and (5) ensures that *PD_u,v_* is set to *D_u_* if *u* is the parent of *v* and that distance of *v* is equal to the distance of its parent *u* increased by 1.

To ensure that 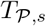 is a tree, we require that each node can have only one parent; this is ensured by constraint (6). We enforce that the nodes of 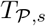 form a connected component of size |*S*| by considering a fictitious *network flow* originating at root node *s* of |*S*| − 1 units – by constraint (7). For each directed edge (*u, v*), the value of flow along that edge is represented by variables *f* (*e_u,v_*).

The flow loses 1 unit at each node - by constraint (8). If there is a positive amount of flow through the edge (*u, v*), then it is a tree edge in 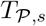 – by constraint (9). If there is no flow through the edge (*u, v*) then it is not in 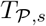 – by constraint (10).

The set of constraints (11–17) ensures that the evolutionary trajectories represented by paths in 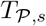 are conserved. Constraint (17) requires each trajectory (or path) from the root *s* to a node *v* in the tree to be conserved in at least 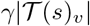 tumor graphs. Similar to the use of auxiliary variables *PD_u,v_* above, we introduce auxiliary variables 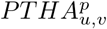, set to 1 if and only if there is a path from *s* to *u* in tumor graph *G_p_*, and (*u, v*) is a tree edge of 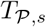 – by constraints (13), (14) and (15). A tree path from *s* to *v* exists in a tumor graph *G_p_* if *G_p_* both includes the path from *s* to *u* as well as the tree edge from *u* to *v* - by constraint (16). Here, the indicator constant *α_u,v_* takes the value 1 if the edge (*u, v*) ∈ *G_p_*, and 0 otherwise. By default, the root node *s* has a path in tumor graph *G_p_* if it is observed in the tumor *p* – by constraints (11) and (12).

Note that only *X_u,v_* variables are required to be binary. The decision variables 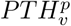 can be continuous, with the addition of constraints that they can not be larger than 1. This reduces the complexity of the model and the running time and space required to solve it.

### 2.2 Not transitive input graphs

Section 2.1 describes finding the evolutionarily conserved alteration set 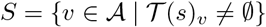 in a prior step to building the maximum conserved evolutionary trajectory tree from a given root *s*. The transitivity of directed edges in the input tumor graphs makes this easy – we simply add to *S* all nodes *v* for which there exists a directed edge (*s, v*) in at least one of the tumor graphs.

However, in some applications the user may wish to break the transitivity in a tumor graph *G_p_* so that while it includes directed edges (*u, w*) and (*w, v*), it does not have (*u, v*). This could be desirable if there is strong belief/confidence that the intermediate node/alteration *w* was necessary for the emergence of alteration *v* in *G_p_*. In such cases, the user may ensure that an evolutionary trajectory involving alterations *u* and *v* does not “skip” a high-confidence intermediate alteration *w* by excluding the edge (*u, v*). Note that this may lead to some nodes being excluded from the resulting consensus tree: e.g. given *u* as the root, suppose that half of the tumor graphs in the cohort include the path *u, w, v* and the other half include the directed edge (*u, v*) but not the directed edge (*w, v*). If *γ* > 0.5 then *v* will not be included in the consensus tree.

Furthermore, the relaxation of the transitivity property makes the problem of finding the evolutionarily conserved alteration set non-trivial; this is due to the fact that in any *G_p_*, not all nodes on the directed path from root *s* to another node *v* will necessarily be the children of *s*. We show how to address this issue by a preprocessing step described here. Let *S*\* be the largest set of nodes such that for each node *v* ∈ *S*\*, there are at least 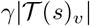 graphs which include a path from *s* to *v* consisting only of nodes in *S*\*. If a node *v* is in the resulting consensus tree, the path from *s* to *v* must consist only of nodes in *S*\* (since *S*\* is defined as the largest such set). This implies that a node that is not in *S*\* cannot be in the consensus tree, implying that computing the set *S*\* and pruning out all nodes not included in it can reduce the solution space. Given a node *s*, we formulate the *maximum path-conserved subgraph identification* problem whose solution is the set *S*\* and describe a polynomial-time algorithm to compute it. Note that on an input set of directed graphs 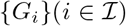, we use the notation 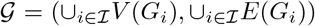 to denote the smallest supergraph of all graphs 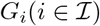 (so that each *G_i_* is a subgraph of 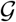).

#### Maximum Path-Conserved Subgraph Identification problem (MPCSI)

Given 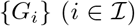, their smallest supergraph 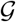, a root node 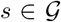, and a number *t_v_* for each 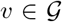, find the largest set of nodes *S*\* of 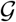 such that for each node *v* ∈ *S*\* there are at least *t_v_* graphs *G_i_* which include a path from *s* to *v* consisting only of nodes in *S*\*.

To solve MPCSI problem for a given root *s*, CONETT uses an iterative algorithm which starts with *S*\* = *s*. In each iteration, the algorithm considers adding to *S*\* a new node *v* to which there is a directed edge from any node in *S*\* in at least *t_v_* graphs. Node *v* is only added to *S*\* if there are at least *t_v_* graphs that include a path from *s* to *v* consisting only of nodes already in *S*\*. For finding out whether *v* satisfies this condition, CONETT constructs a data structure *P* (*v*) for each node 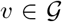, which maintains the set of graphs that have a directed edge from any node in *S*\* to *v* in the form of a bitmap. Naturally *P* (*s*) is initially equal to the set of all graphs where *s* is present. For every other node *v*, *P* (*v*) is initially equal to the set of tumor graphs that include edge (*s, v*). The algorithm terminates if there is no 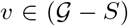 such that |*P_v_*| ≥ *t_v_*.

Otherwise it identifies any node 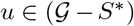 such that |*P_u_*| ≥ *t_v_* and adds it to *S*\*. Then it checks for each node *v* ∈ *S*\* for which 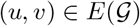, whether the addition of *u* to *S*\* adds any new graphs to *P_v_*. If this is the case, the algorithm inserts the new graphs to *P_v_* and recursively propagates the new graphs to any neighbour *w* ∈ *S*\* of *v* that does not already have them in *P_w_*. Figure S-1 illustrates the necessity of the recursive path propagation within *S*\*. Finally, the algorithm updates *P_z_* for every node 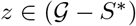 for which there is an edge 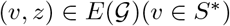.

The correctness of the algorithm follows from the invariant that at each iteration, the algorithm considers paths from *s* to nodes that are either in *S*\* or are neighbours of nodes in *S*\*, thereby ensuring the existence of at least *t_v_* paths (each in a separate graph). We also note that for any given *v* ∈ *S*\*, the total number of times that *P_v_* is updated by the algorithm can not be more than the number of graphs. Furthermore the total number of times the algorithm can visit *v* is no more than the number of edges in 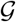. As a result, the running time of the algorithm is 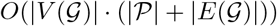.

### 2.3 Optional Constraints and Parameters

CONETT allows the user to add additional constraints to the ILP solution to the *MCETT* problem described in Section 2.1, via a new set of options/parameters as follows: (1) CONETT allows the user to restrict the set of nodes which are included into the consensus tree to more recurrent alteration events that “follow” the root event in a specified minimum number of tumor graphs *t*. This is achieved by removing all nodes *v* from the evolutionarily conserved alteration set *S* for which 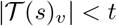, prior to building the tree.

(2) CONETT also allows the user to add constraints on the ancestor-descendant ordering of nodes in conserved evolutionary trajectories of the resulting tree 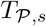. Specifically, it calculates a “confidence score” of *u* being an ancestor of *v* based on evidence provided by all tumor graphs in which root *s* is an ancestor of *u* and *v* that do not have “don’t care” edges. Given a user-defined value *δ*, representing the threshold for confidence scores between each pair of nodes in the evolutionarily conserved alteration set *S*, CONETT adds constraints to the ILP formulation that require each ancestor-descendant pair of nodes in the tree to have a confidence score at least *δ*.

CONETT’s confidence score for node *u* being an ancestor of node *v*, penalizes those tumor graphs that include (i) a directed edge (*v, u*) without a directed edge (*u, v*) and (ii) an anti-edge between *u* and *v*, while considering tumor graphs with a don’t care edge between *u* and *v* neutral. Specifically, let 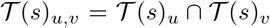 be the subset of all tumor graphs in which there exists a directed path from *s* to both *u* and *v* (note that no tumor graph in this set has a “don’t care” edge between *u* and *v*). Let 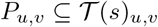 be the subset of tumor graphs that contain a directed edge (*u, v*) (the presence of directed edge (*v, u*) does not make a difference). Also let 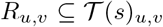 be the set of tumor graphs that include the directed edge (*v, u*) but not the edge (*u, v*). Similarly, let 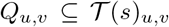 be the subset of tumor graphs which include an anti-edge between *u* and *v* ^4^. Note that 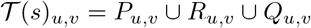. Then the confidence score of *u* being and ancestor of *v* in the subset of tumor graphs 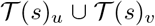 is set to be 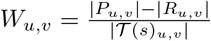. As can be seen below, this (as well as alternative) confidence score(s) penalizing (or incentivizing) specific edge types is easily incorporated in CONETT’s ILP formulation.

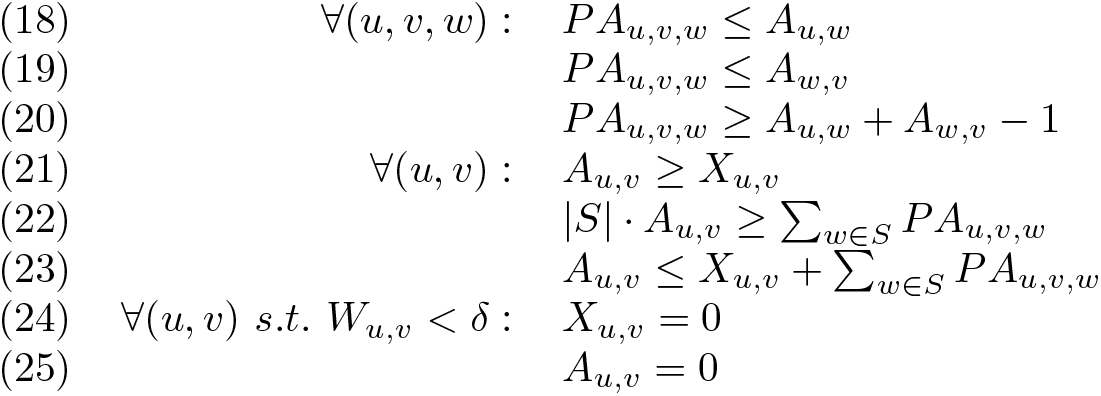

For each pair of nodes *u* and *v* in *S*, we add a variable *A_u,v_*, which is set to 1 if *u* is an ancestor of *v* in 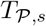, and 0 otherwise. Constraints (24) and (25) ensure that all ancestor-descendant pairs of nodes in 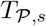 have a high confidence score. We introduce auxiliary variables *PA_u,v,w_*, set to 1 if *u* is an ancestor of *w* and *w* is an ancestor of *v* and set to 0 otherwise – by constraints (18), (19) and (20). A parent of a node is its ancestor by default, by constraint (21). Constraints (22) and (23) impose transitivity on ancestor-descendant relationships: *u* is an ancestor of *v* if and only if *u* is an ancestor of *w* and *w* is an ancestor of *v*, for some intermediate node *w*.

### 2.4 Assessment of the statistical significance of MPCSI solutions

In order to assess the statistical significance of an evolutionarily conserved alteration set *S* computed by CONETT (or a solution to the MPCSI problem for graphs that are not transitive), we perform a variation of the standard permutation test on the input data. Let *N_a,p_* represent the number of nodes (genes and chromosome arms) altered by alteration type *a* in tumor graph *G_p_*, and let *F_v,a_* represent the number of tumors in which node *v* occurs altered by *a*. ^5^ For each tumor 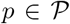, and each alteration type *a*, we randomly draw (without repetitions) *N_a,p_* nodes, and mark them as altered by *a*, in case they were altered by *a* in at least one tumor graph in the original input. The nodes are randomly drawn from a distribution based on their recurrence frequency, i.e. each node *v* is drawn with the probability 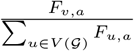. That way, the average recurrence frequency of each node and alteration type in the permuted data is equal to the original recurrence frequency, preserving hidden inter-dependencies among somatic mutations and copy number alterations that were the cause of the observed recurrence frequencies and evolutionarily conserved alteration sets. Additionally, consistent with observations on the TRACERx ccRCC data set we use in this paper, special care was devoted to ensuring that a copy number event affecting a whole chromosome does not appear together with a copy number event altering either of its arms within the same tumor. Lastly, the randomly chosen altered nodes replace the original nodes of the input tumor graph *G_p_* after being subject to a random permutation.

The above process is repeated 1000 times. In order to estimate the statistical significance of a set *S* of size *k*, obtained with root alteration event *s* that occurs altered in *F_s_* tumors, we run the method on each generated permutation of the input data using the same parameters, and record the number of times that the resulting set has a size of *k* or greater when computed with a root that occurs altered in *F_s_* or more tumors. That number, divided by the total number of experiments (1000 for our case), gives us the empirical p-value estimate for *S*.

## 3 Results

### 3.1 Data and pre-processing

#### 3.1.1 Clear-cell renal cell carcinoma (ccRCC)

We obtained ccRCC data from the TRACERx renal study [31], which gathered multi-region sequencing data from 100 tumors. The study collected non-silent mutation data in about 110 genes that are deemed to be high-confidence ccRCC driver genes, obtained through a sequencing panel from which single nucleotide variants (SNVs), dinucleotide variants (DNVs) and small insertion and deletions (INDELs) were inferred. The study also detected somatic copy number alterations (SCNAs), which were reported only on the level of chromosome arms or whole chromosome – provided at least 50% of the chromosomal arm had a copy number alteration. Then, in each tumor, the study clustered all the alterations and reconstructed a phylogenetic tree of clonal hierarchy.

For every somatically altered gene and copy number altered chromosomal arm that is found in a particular tumor’s phylogenetic tree, we created a node labeled by the name of the gene (or chromosome) and the type of alteration that affects it. Since, in this dataset, SNVs, DNVs and small insertion and deletion events result in inactivation of the genes that they affect, we treated them all as a single “inactivating” alteration type. Then, we build a directed graph on this set of nodes for a particular tumor, such that a directed edge is drawn from node *u* to node *v* if the clonal node in the phylogenetic tree which carries the alteration affecting node *u* is an ancestor of the clonal node in the phylogenetic tree which carries the alteration affecting node *v*. In the case of alteration events *A* and *B* that belong to the same clone, we draw parallel directed edge, in opposite directions, between the nodes that are associated with *A* and *B*. In 5 tumor trees, there was immediate branching right after the germline node. To ensure that there are no cross links in our tumor graphs (which can happen if a gene is present in both branches), leading to false precedence relationship between ancestors of the cross link source and descendants of the cross link destination, we split such trees at the root and treat them as two separate tumor phylogenetic trees.

As a result, we obtained tumor graphs from 105 “tumors” (after splitting the 5 trees with branching at germline node), containing 242 different nodes in total, representing driver genes affected by inactivating sequence alterations, and copy number gains and losses on chromosomes – both on whole chromosome and arm level. Figure S-2 shows recurrence frequencies of the 242 events; the *y* axis shows the recurrence frequency (in log scale), and events are sorted along the *x* axis based on their *y* values for clarity purposes. The figure reveals the sparsity of the data – more than a third of genes that carry sequence alterations each occur in only a single tumor, and more than a half occur in at most 2. Among more recurrently altered genes and chromosome arms (those that are found altered by the same alteration type in at least 10 tumors), there are 19 (35%) copy number gain-altered, 26 (50%) copy number loss-altered and 12 (9%) sequence-altered nodes.

#### 3.1.2 Non-small-cell lung cancer (NSCLC)

We obtained data from 99 NSCLC patients from [5]. Originating from whole-exome sequencing of multiple spatially separated regions from the TRACERx lung cancer study [18], [5] provide cancer cell fraction (CCF) values, as well as clustering information, for single nucleotide variations and focal copy number alterations in 79 putative driver genes of NSCLC (we note that there was no case of a gene being affected by alterations of different types in any patient). Using CITUP [24], we inferred phylogenetic trees with the clustering information provided by [5], and for each patient we selected the tree with the minimum error score (which was always unique). We then construct tumor graphs from the phylogenetic trees similarly to the procedure used to generate tumor graphs in the ccRCC data (Section 3.1.1).

Contrary to the TRACERx ccRCC data, in which the clustering of alterations contained chromosome-level copy number alterations, the clustering of alterations did not contain copy number alterations on a chromosome level in this dataset. Since in [5], the analysis was performed on the gene level, ignoring the specific alteration type (or having just one universal alteration type), we took a similar approach to make our results comparable by merging SNVs and gene deletions into a single “inactivating” alteration type. However, we kept gene amplifications as separate alteration type since the downstream effect significantly differs from SNVs and deletions.

### 3.2 Conserved evolutionary trajectories in ccRCC

We first used CONETT to identify statistically significant alteration sets that consisted of highly-recurrent alteration events following the root event, and associated trajectories in the TRACERx renal study (see Section 3.1.1), by considering as a root node each gene and chromosomal arm that are found to be altered in at least 10 tumors. For each root, in this experiment we set the optional parameter *t* in the MPCSI formulation to a high fraction of the root’s recurrence frequency (≥ 48%). We empirically assessed the statistical significance of these alteration sets using methodology described in Section 2.4. From the evolutionarily conserved alteration sets, we constructed conserved evolutionary trajectory trees, setting the *γ* parameter in the *MCETT* formulation to 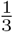 and the ancestor-descendant order confidence score threshold of *δ* = 0.85 (see Section 2.3). Below we report on the 10 most significant evolutionarily conserved alteration sets.

Four of these sets contain 2 or 3 nodes, involving genes and copy number altered chromosome arms that are co-clonal (Figure 3E): these include sequence alterations on driver gene VHL and copy number loss on chromosome 3p, which are known to be founder alterations in ccRCC. Conserved evolutionary trajectories rooted either by sequence alterations on VHL or copy number loss on 3p (Figure 3A-B) both include sequence alterations in PBRM1 and copy number loss on chromosome 14q as highly-conserved follow-up events (p-value < 0.001), with sequence alteration in PBRM1 typically occurring before copy number loss on 14q. Copy number gain on chromosome 5 (occurring in about a quarter of all tumors) is accompanied by co-clonal sequence alteration in VHL, and is followed by copy number loss on chromosome 14q and copy number gain on chromosome 7, each observed in about half of the tumors (p-value < 0.008, Figure 3C). The remaining three conserved trajectories are all rooted by the copy number loss on chromosome 6, occuring in 19 tumors (Figure 3D shows the largest trajectory tree). It is followed by losses on 14q and 9 at a high conservation rate of 70% (p-value < 0.001), gains on chromosome 7 at 55% (p-value < 0.010) and sequence alterations of VHL and loss on chromosome 4 at 50% (p-value < 0.001). Loss on chromosome 4 follows loss of chromosome 9 in more than a third of the tumors.

**Figure 3:**
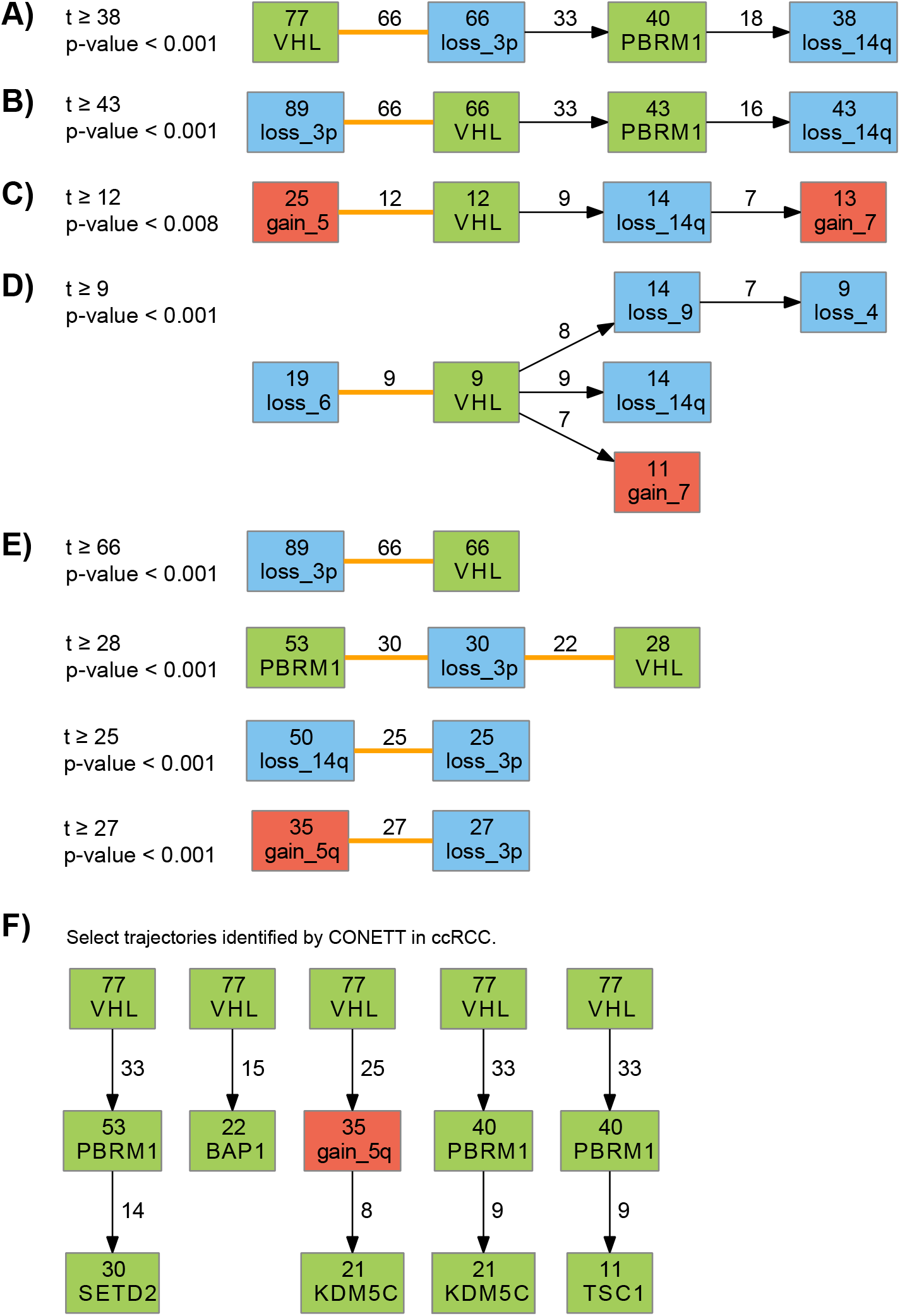
Conserved evolutionary trajectories identified through CONETT in renal cancer. The numerical value in each node is the total number of tumors in which the root event is an ancestor of the node. Next to each edge, the number of tumors that the exact evolutionary trajectory ending with that edge is conserved in is displayed. Each undirected edge (orange) marks a pair of co-clonal alterations that can not be temporally ordered. **A, B, C, D**) Conserved evolutionary trajectory trees constructed for highly-conserved alteration sets. **E**) Statistically significant highly-conserved, co-clonal sets, from which it is not possible to reconstruct evolutionary trajectories with a known temporal order of nodes. **F**) Select trajectories from conserved evolutionary trajectory trees rooted by the germline node, *VHL* and loss on chromosome 3p.

Next, we looked into trees of trajectories rooted by the most recurrent events in the data set. As sequence alterations in VHL (found in 77% tumors) and copy number loss on chromosome 3p (found in 89% tumors) play a key role in initiation of ccRCC, we examined the two evolutionarily conserved alteration sets and conserved evolutionary trajectories that are rooted by either of these events and contain nodes that are altered in at least 10 tumors each (*t* = 10, both of the sets have p-value < 0.001). We also examined conserved evolutionary trajectories that start with the germline node, with the same value of *t*, as it is present in the whole data set. The resulting trees of trajectories are shown on Figures S-6, S-7, and S-8; while Figure 3F shows a number of select trajectories from those trees.

Consistent with the trees reported in the original TRACERx renal study, the trajectories obtained by CONETT include a series of sequence alteration events *VHL* → *PBRM1* → *SETD2* observed in 14 tumors, as well as *VHL* → *BAP1* observed in 15 tumors. Additionally, CONETT also detected two conserved evolutionary trajectories terminating with a sequence alteration in gene KDM5C that collectively cover 80% of tumors where it is present: *VHL* & *loss_3p* → *gain_5q* → *KDM5C* in 8 tumors and *VHL* → *PBRM1* → *KDM5C* in 9 tumors; these trajectories were not reported in the original study. Note that KDM5C is a chromatin modifier, similar to BAP1; its dysregulation alters the activity of many other genes [13]. Additionally, *VHL* → *PBRM1* → *TSC1* is found in 9 tumors, representing 75% tumors where *TSC1* is present. CONETT also identified a number of less frequently altered genes that do not display high confidence ancestor-descendant ordering. These alteration events in genes such as *MTOR* (in 12/17 tumors), *OBSCN* (in 7/12 tumors), and *MUC16* (in 10/16 tumors) all seem to branch out from an alteration in *VHL*.

Inactivations of both *PBRM1* and *BAP1* are known to rarely co-occur in ccRCC [28, 31], and trees of conserved evolutionary trajectories on Figures S-6, S-7 and S-8 always place them into different branches. However, most of the time when they do co-occur, *PBRM1* alteration precedes *BAP1* alteration. We thus ran CONETT using *PBRM1* alteration as the root event, to find out whether BAP1 sequence alteration appears as a conserved folow-up event, using *t* = 10 (p-value < 0.001). The resulting tree of conserved evolutionary trajectories is shown on Figure S-9. Alteration on *BAP1* was not identified as a downstream event, strengthening the hypothesis that their co-occurrence is most likely due to upstream events such as loss on 3p and inactivation of *VHL*.

Loss on chromosome 14q is the second most recurrent large-scale copy number loss event (after loss on 3p), occurring in 50 tumors. However, contrary to loss on 3p which regularly appears as the earliest alteration event in tumors, loss on 14q is always placed later, after both VHL and loss on 3p. It is often preceded by inactivation of *PBRM1*, although not in every single tumor (Figure S-9 shows that loss on 14q follows inactivation of *PBRM1* in 22 tumors). There is also strong evidence of precedence of gains on chromosomes 5 and 5q (which are themselves mutually exclusive) to loss on chromosome 14q – trajectories rooted by loss on 14q do not detect gains on chromosome 5 or 5q as downstream events (Figure S-10), but trajectories rooted by gains on 5 and 5q do detect loss on 14q as a downstream event (Figure S-11). Additionally, Figure S-11A shows presence of mutual exclusivity of losses on chromosome 3p and 14q in tumors that contain gain on chromosome 5, whereas that relationship is not noticed in tumors that contain gain on 5q (Figure S-11B). This pattern of mutual exclusivity was not reported in the original ccRCC TRACERx study.

### 3.3 Conserved evolutionary trajectories in NSCLC

We used CONETT to identify conserved evolutionary trajectories in the TRACERx non-small-cell lung cancer study (see Section 3.1.2). This data was analyzed by REVOLVER [5], allowing us to compare the CONETT-identified trajectories with the edges in clusters identified by their method. Since REVOLVER uses “collective” information from the whole set of tumors to infer individual tumor phylogenetic trees, in order to make our results more comparable we set the value for the optional parameter *δ* = 1 – ensuring that the temporal order of ancestor-descendant pairs of nodes in our trees is conflict-free with not just those tumors in which the specific trajectory that they lie on is conserved, but also the wider set of tumors in which the root is ancestor of both the nodes (please see Section 2.3 for exact details). We used the same value for the parameter 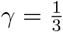 as in the analysis of the ccRCC dataset.

In [5], the authors highlighted evolutionary trajectories constructed from edges in cluster C5 (Figure 4B). Using CONETT, with the SNV-altered gene *CDKN2A* as the root, we were able to recover the full topological order of the genes in cluster C5 (Figure 4A), as well as additional ancestor-descendant relationship between amplification on *TERT* and SNVs on *FAT1* and *NOTCH1*. The CONETT-identified tree captures more occurrences of tree edges *CDKN2A* → *TP53* and *TP53* → *TERT*, as well as edges from *TP53* towards *NOTCH1* and *COL5A2*, which were tree edges in cluster C5 but are just forward edges in CONETT (dashed gray lines); due to CONETT trajectories capturing additional ancestor-descendant relationship between amplification of *TERT* and SNVs on sibling nodes *FAT1* and *NOTCH1*.

**Figure 4:**
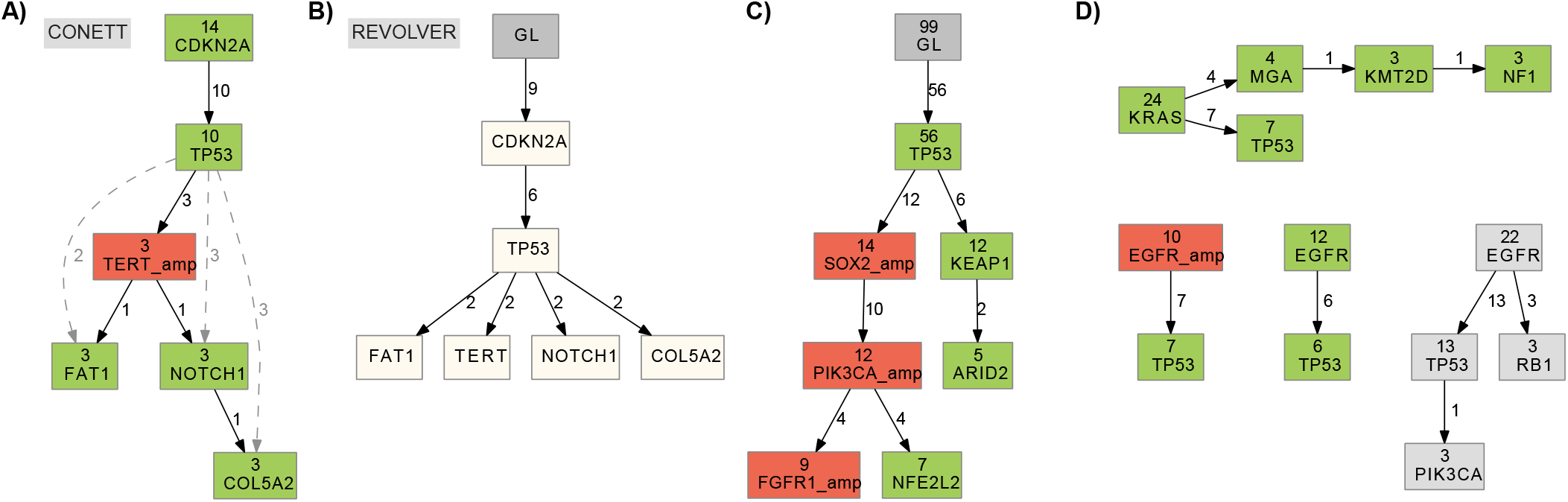
Conserved evolutionary trajectories identified through CONETT in lung cancer. The numerical value in each node is the total number of tumors in which the root event is an ancestor of the node. Next to each edge, the number of tumors that the exact evolutionary trajectory ending with that edge is conserved in is displayed. **A)** CONETT-constructed tree of conserved evolutionary trajectories rooted by an SNV on the gene *CDKN2A* (*t* = 3). It captures all the genes shown in the likely tree of cluster C5, selected by REVOLVER (shown in B). It captures more occurences of the genes and the edges and it shows additional ordering of the alterations. **B)** A likely evolutionary tree drawn from the edges in cluster C5, identified by REVOLVER. Edge labels show the number of occurrences of the edge within the cluster. **C)** Select trajectories identified by CONETT which have a *TP53* SNV as the clonal event. **D)** Trajectories rooted by recurrent NSCLC drivers *KRAS* and *EGFR*. Rightmost tree has EGFR amplifications and SNVs merged together.

Additionally, we used CONETT to construct trees of trajectories with known early drivers (as well as the germline node) as roots (Figures S-12, S-13). Figure 4C shows select conserved trajectories which are supported by both trees. The trajectories capture *TP53* → *SOX2* & *PIK3CA* amplifications → *NFE2L2* (in 4/9 tumors), as well as *TP53* → *KEAP1* (in 6/12 tumors) – inactivations of *NFE2L2* and *KEAP1* may be linked to increased chemoresistance [19]. All *TP53*-rooted evolutionarily conserved alteration sets were found to be statistically significant (p-value < 0.001). Interestingly, trajectories rooted by SNV and amplification of *EGFR* did not pick up any node other than *TP53*, which they are often found by REVOLVER to be in the same cluster with (Figure 4D). Even when we merge amplifications and SNVs of *EGFR*, *RB1* and *PIK3CA* just barely emerge as downstream altered genes. *KRAS*-rooted trajectories capture *MGA* and *KMT2D*, consistent with the germline-rooted tree (Figure S-12).

## 4 Discussion

We present CONETT, a novel computational method for combinatorial detection of conserved evolutionary trajectories in tumors. CONETT is the first such method that goes beyond just identifying clusters of common edges and directly produces a consensus phylogeny which aims to maximize the number of ancestor-descendant ordered pairs of alteration events. Given a root alteration event, we show how to construct a conserved evolutionary trajectory tree consisting of the maximum set of events that can be found on conserved evolutionary trajectories from the root via an ILP formulation. We applied our method to recently published datasets of 100 ccRCC tumors and 99 NSCLC tumors, and identified conserved evolutionary trajectories of sequence-altered genes and copy number altered chromosomal arms. CONETT identified all conserved evolutionary trajectories involving sequence altered genes reported as significant by the original studies; it also identified several additional conserved trajectories which were not reported earlier.

As per the original studies of TRACERx renal and lung cancer data, we have not differentiated sequence-alteration types and have treated all sequence-altered genes identically; this allowed us to compare CONETT results with those of the original studies. However CONETT’s framework is sufficiently general to offer the user the ability to differentiate somatic alteration types in the trajectories it identifies so as to perform in depth analysis of larger and richer data sets to be published in the near future.

We acknowledge that the consensus tree that CONETT’s ILP formulation currently constructs represents a single tree topology in which each node occurs only once. That forces the method to select only a single trajectory from a given root towards each node, even though there might be multiple different trajectories that each occur in a significant fraction of tumors (as evidenced by the two different trajectories towards *KDM5C* discovered by CONETT and shown on Figures S-6 and S-8, together covering 80% of tumors in which the gene is sequence-altered). In future, it would be useful to devise a way to simultaneously identify different conserved trajectories in tumor evolution towards the same node.

As larger and more precise data sets emerge, as well as depending on the motivation for and the specific application of the analysis, it might become desirable to use an alternative version of tumor graphs which do not have the property of transitivity of the directed edges. This could be desirable if there is strong belief that an ancestor event *u* was necessary for the emergence of another follow-up event *v*. In such cases, removing transitive edges towards *v* would ensure that detected evolutionary trajectories involving *v* would not be able to “skip” the high-confidence intermediate alteration event *u*. Our method can incorporate such data into its framework as is; we also give a polynomial time algorithm to detect a set of events that is a superset of the nodes in the resulting tree – allowing to prune the search space and reduce the running time and space of the ILP model.

Additionally, we introduced an optional scoring scheme which measures the confidence of ancestry for a pair of nodes based on the evidence given by the temporal order of the two events in the input tumor data. Currently, our method uses a common threshold for all possible ancestor-descendant pairs of nodes, imposing a uniform level of stringency. It is worth exploring how to impose a level of stringency specific to each individual pair of nodes on evidence available for their partial ordering in the future.

## Supplementary Materials

### 1 Supplementary Methods

#### 1.1 Polynomial-Time Algorithm for the Maximum Depth Spanning Tree Problem

##### Maximum depth spanning tree (MDST)

Given an unweighted, directed graph *S* and a root node *s* ∈ *S*, find a spanning tree with the maximum total node depth.

We solve the MDST problem via an incremental algorithm which starts with the node *s* and then expands the tree, one node at a time. We use a modified version of depth-first search (DFS) to add new nodes that do not already belong to the tree, and then check whether depth of any nodes that are already in the tree can be increased via the new node (algorithms 1 and 2).

###### Lemma 1. Algorithm 1 maximizes the total node depth.

*Proof.* Let *u* be a newly added node at the current iteration of the DFS recursion. First, we show that in the current iteration node *u* has no better choice for a parent in the tree than the one that added it. If it did, then there would be an edge from another node in the tree towards *u*, and by nature of DFS, it would have already been added to the tree before. Hence this must be the first time that we have encountered node *u* and the current placement is the only possible one.

Next, we show that edge choices that the algorithm picks are optimal by proving the correctness of the update procedure. The proof follows via the DFS tree created by the order in which nodes are visited. Let (*u, v*) be an edge considered by the algorithm. Note that forward and back edges can be safely ignored. If (*u, v*) is a a forward edge, then *v* has already been processed by the recursion and *D_v_* ≥ *D_u_* + 1, so no change is necessary. If (*u, v*) is a back edge, then *v* is an ancestor of *u*, and including this edge would create a cycle. In order to break the cycle, some node *w* on path *v* → … → *u* would have to be unlinked from its current parent and become child of an ancestor of *v* in order to keep the graph connected. Since any ancestor of *v* has depth that is smaller than depth of *v* by at least 1, this edge swap at best keeps the depth of all nodes in the cycle the same as before. Let (*u, v*) be a cross edge. If *D_v_* ≥ *D_u_* + 1, then the depth of node *v* and its descendants would not increase by changing its parent to *u*. If *D_v_* < *D_u_* + 1, then by unlinking *v* from its current parent and placing it as a child of *u*, algorithm 2 would increase depths of *v* and all its descendants. However, since there might be an ancestor of *v* that *u* also has an edge towards, we do not want to connect *u* to *v*, but to the ancestor. Since this is true for every such ancestor, it follows that updating nodes in topologically sorted order results in optimal depth.

From the previous two paragraphs, it follows that at every iteration of the algorithm, the extension step correctly places node *u* in the tree, and correctly updates all remaining nodes’ parent edges. Therefore, at the last iteration, we will have an optimal tree.

###### Lemma 2. Running time of algorithm 1 on a connected graph G = (V, E) is O(|V||E|).

*Proof.* The main DFS skeleton of the code traverses the graph in a depth-first manner, adding each node that is already not in the tree in total time *O*(|*E*| + *f* (*n*) · |*V*|), where *f* (*n*) is the time needed to update the tree after each newly added node. In the worst case, the new node can cause depth update for each node in the tree. Each update needs to be pushed from the updated node down to its whole subtree. In order to avoid updating same nodes multiple times, we topologically sort the nodes and push updates starting with nodes that are closest to the root and doing the same for every subsequent node whose depth isn’t already updated by one of the previous nodes. Topological sort is carried out in time *O*(|*V*|) via counting sort, since we already have depth values of the nodes and they are integers smaller than |*V*|. Since each node gets visited and updated at most once via normal BFS, total time that the update step takes is *O*(|*V*| + |*E*|). Since our graph is always connected, i.e. |*E*| = Ω(|*V*|), the total running time of the algorithm is *O*(|*V*||*E*|).

**Algorithm 1:**
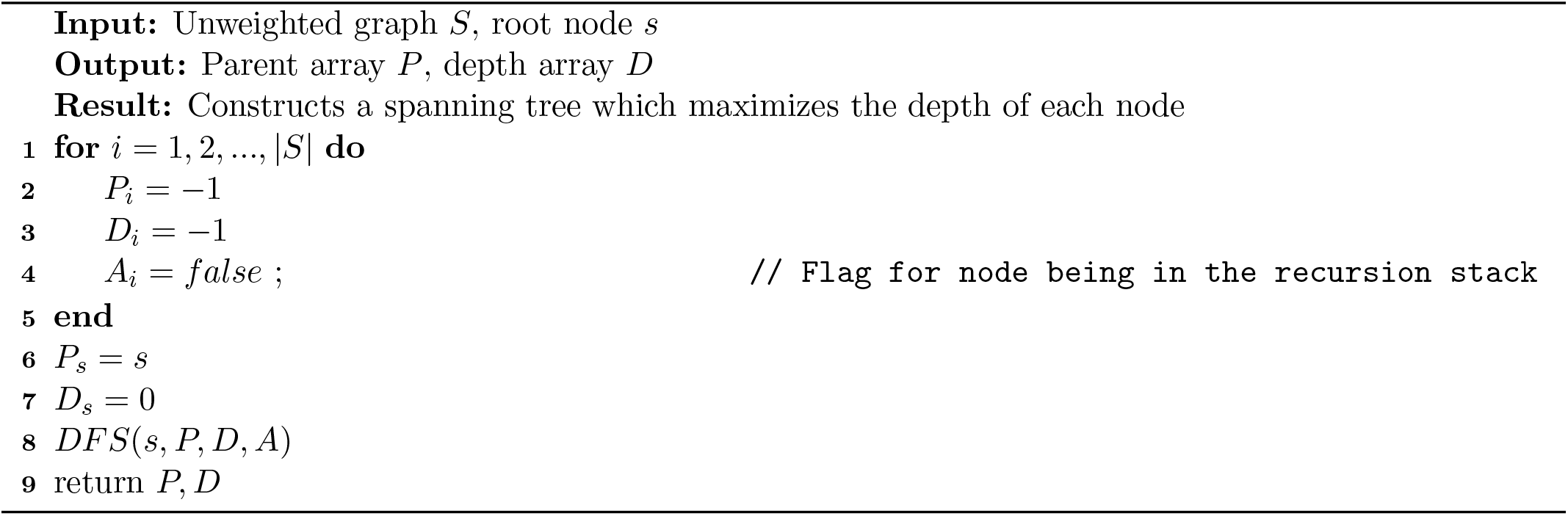
Maximum Depth Spanning Tree

**Algorithm 2:**
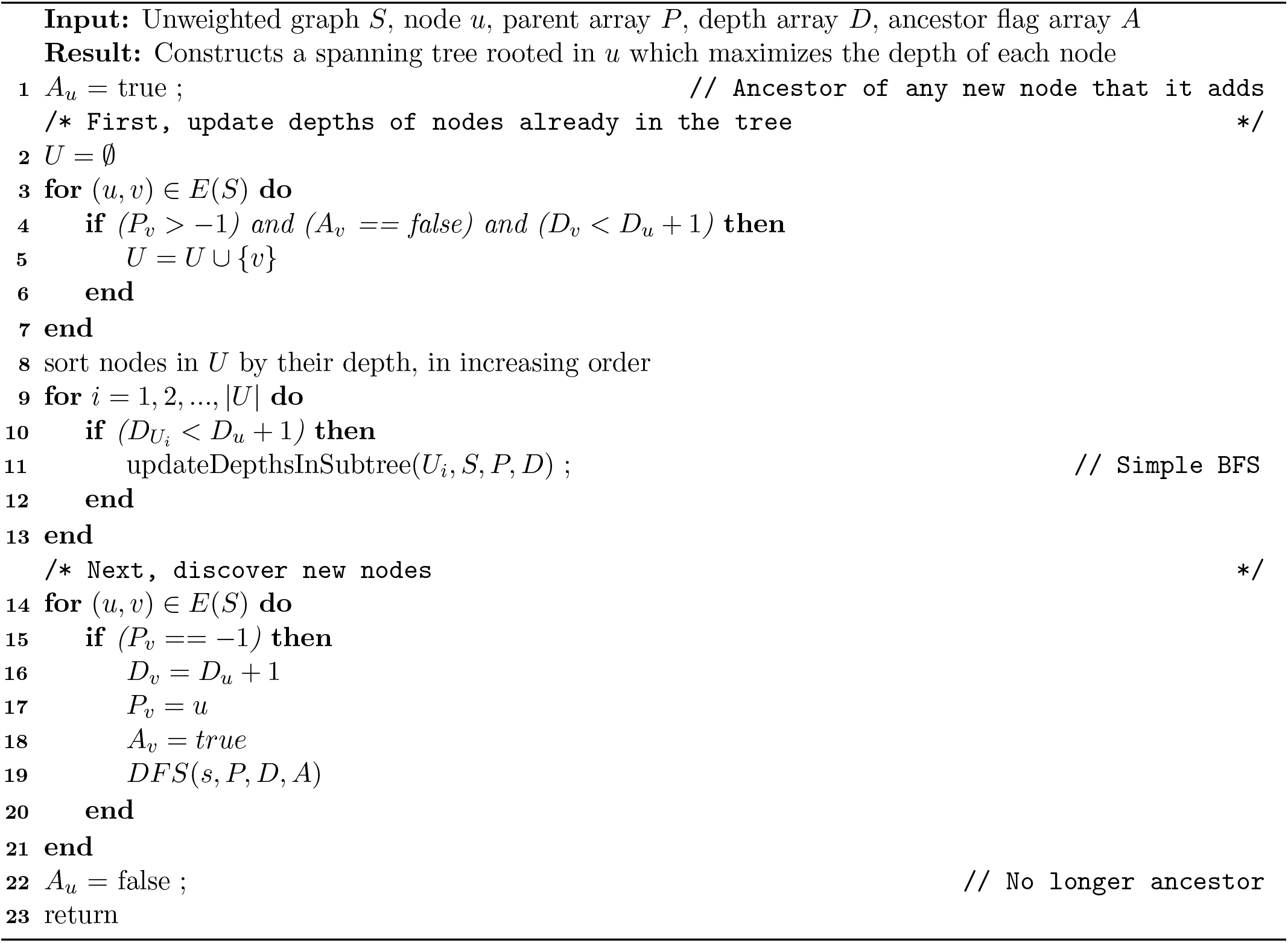
DFS(u, S, P, D, A)

**Algorithm 3:**
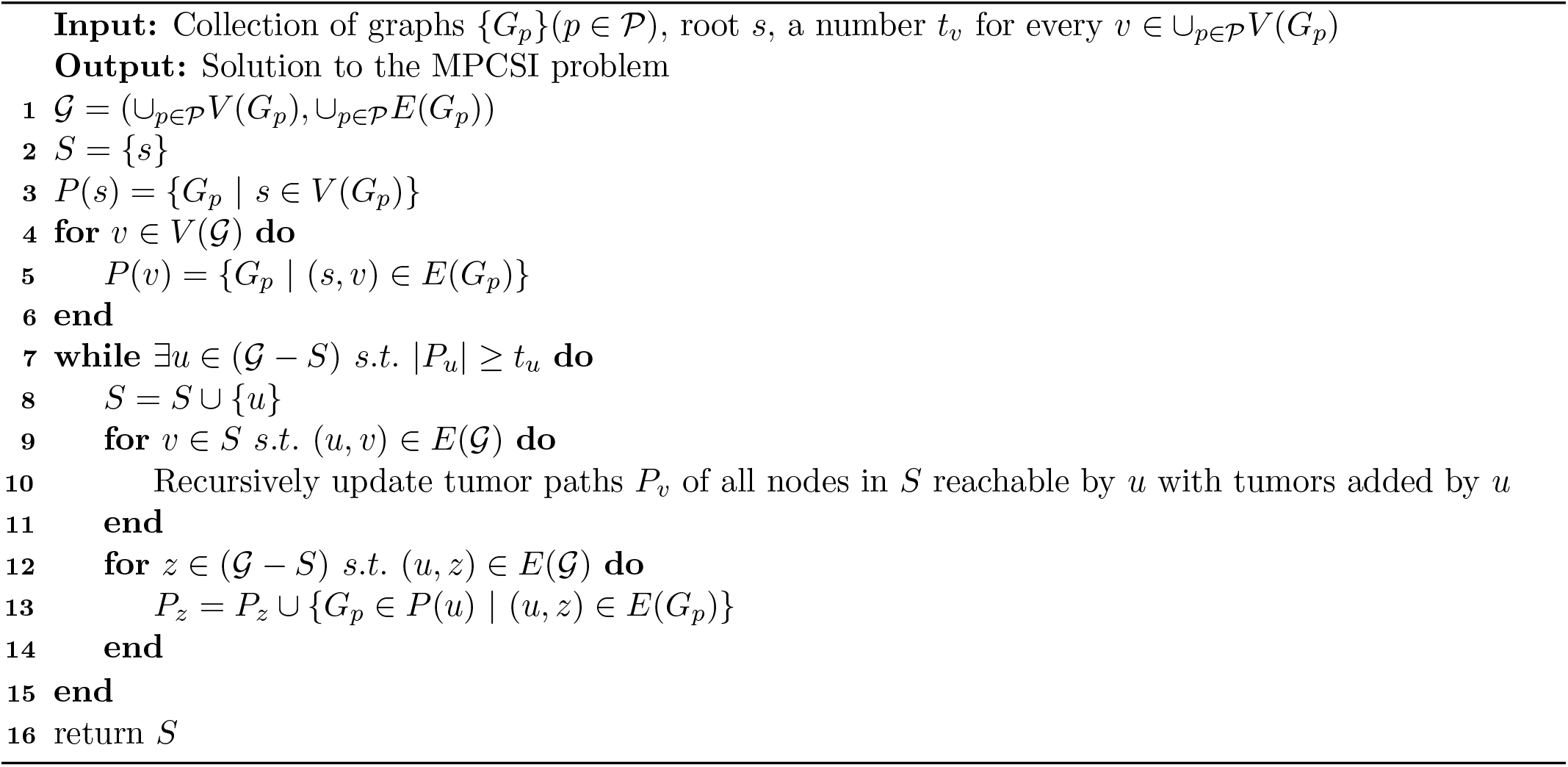
MPCSI

### 2 Supplementary Figures

**Figure S-1:**
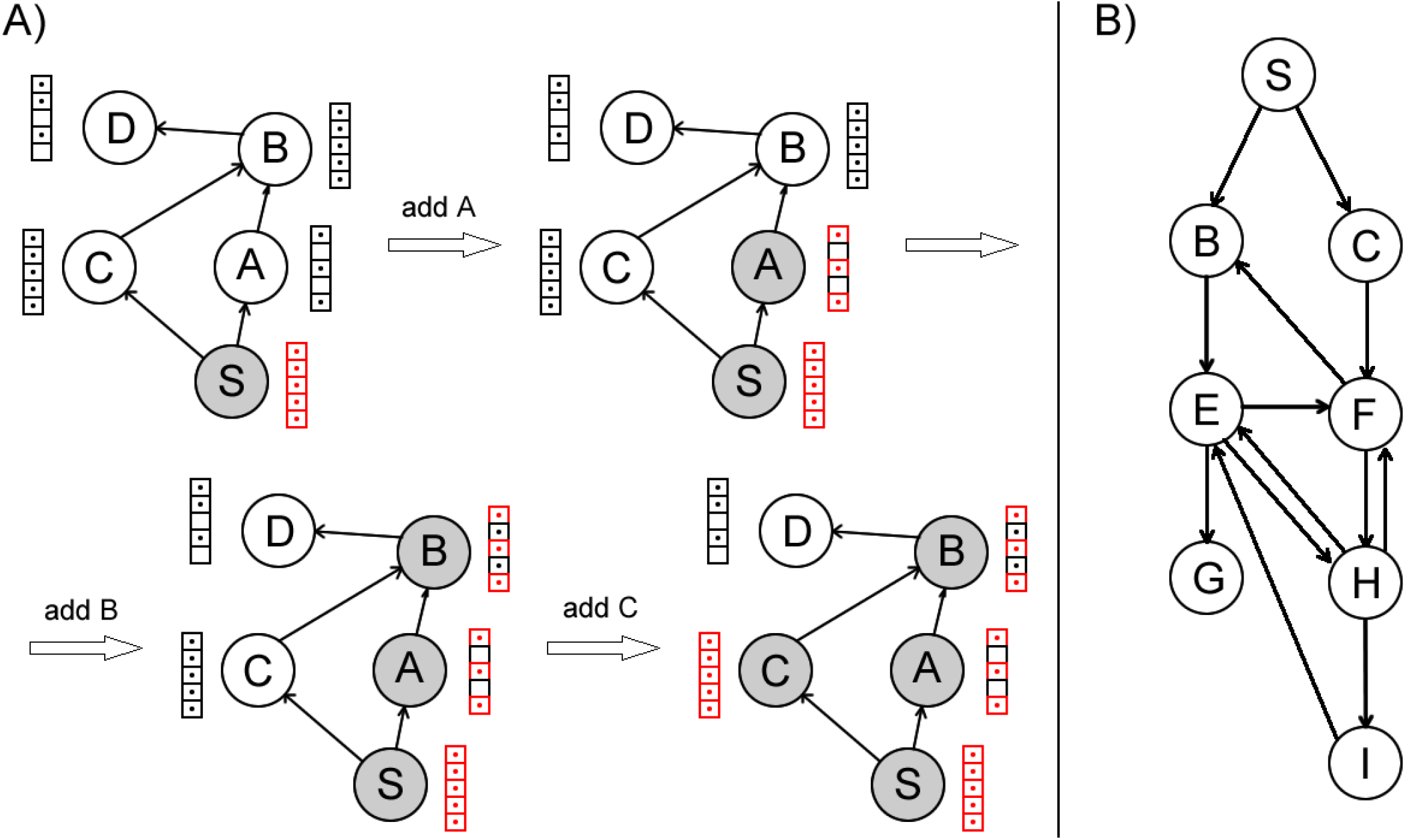
Illustration of the importance of propagation of new paths throughout the subgraph during execution of the MPCSI-solving algorithm. Boxes next to each node represent tumors (total of 5), and there is a dot in a box if the corresponding tumor contains the alteration in the node. **A)** Let *t* = 3. If we add nodes to *S* in order *S* → *A* → *B* → *C*, then *D* cannot be added unless *C* updates paths of *B*. Since the graph has a topological ordering, adding nodes in topological order would avoid the need for updating subgraph nodes’ paths, e.g. *S* → *A* → *C* → *B* → *D*. **B**) This graph contains multiple cycles and does not have a topological ordering of its nodes. Each newly added node has to update paths of every other subgraph node before they are pushed to outside neighbours of the subgraph.

**Figure S-2:**
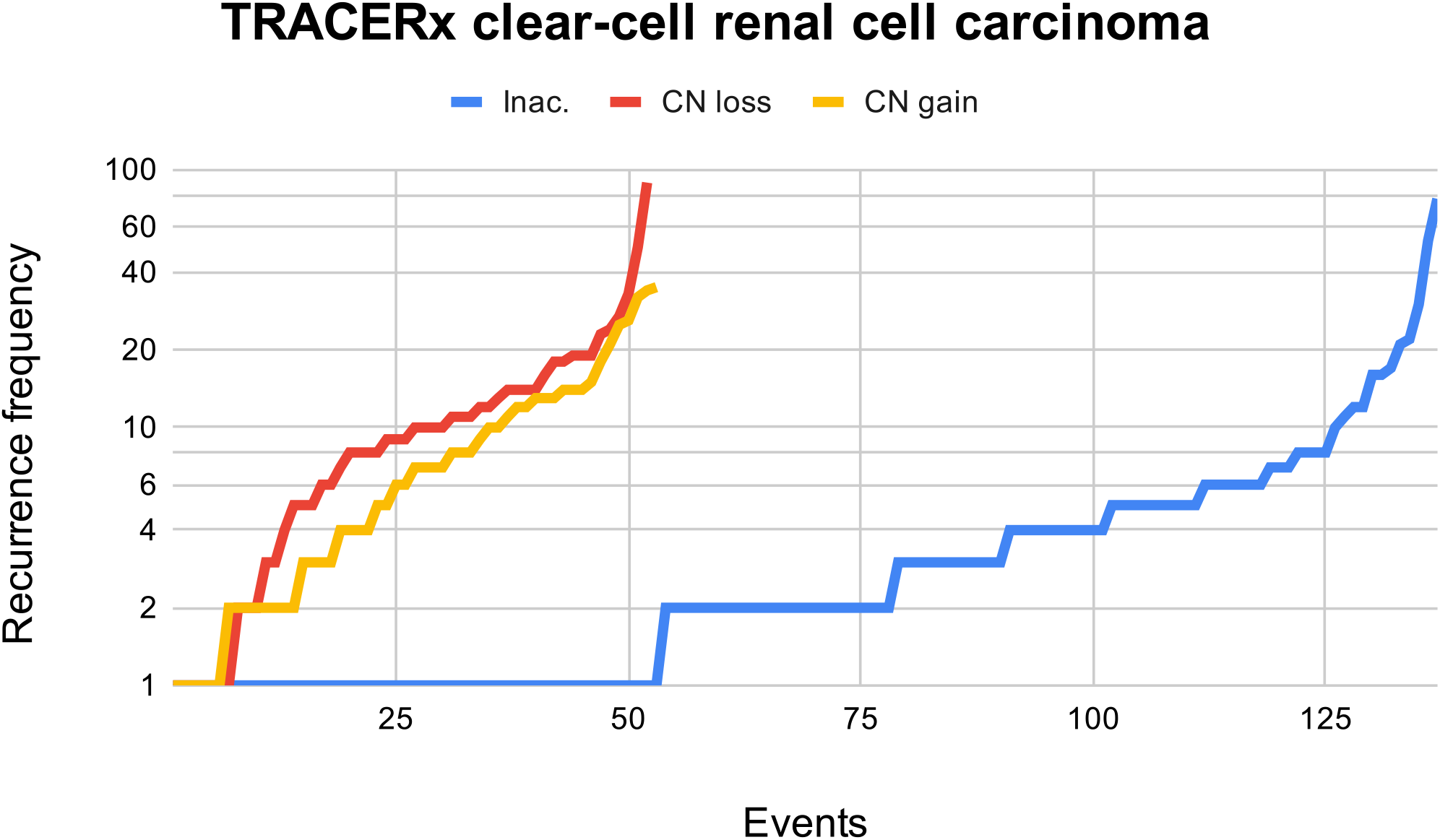
Recurrence frequency of altered genes and chromosome arms in TRACERx renal cohort. The plot shows recurrence frequency of 137 inactivating sequence alteration (blue), 52 copy number loss (red) and 53 copy number gain (yellow) events. Events are sorted by their *y* value along the *x* axis for clarity. The *y* axis is in log scale.

**Figure S-3:**
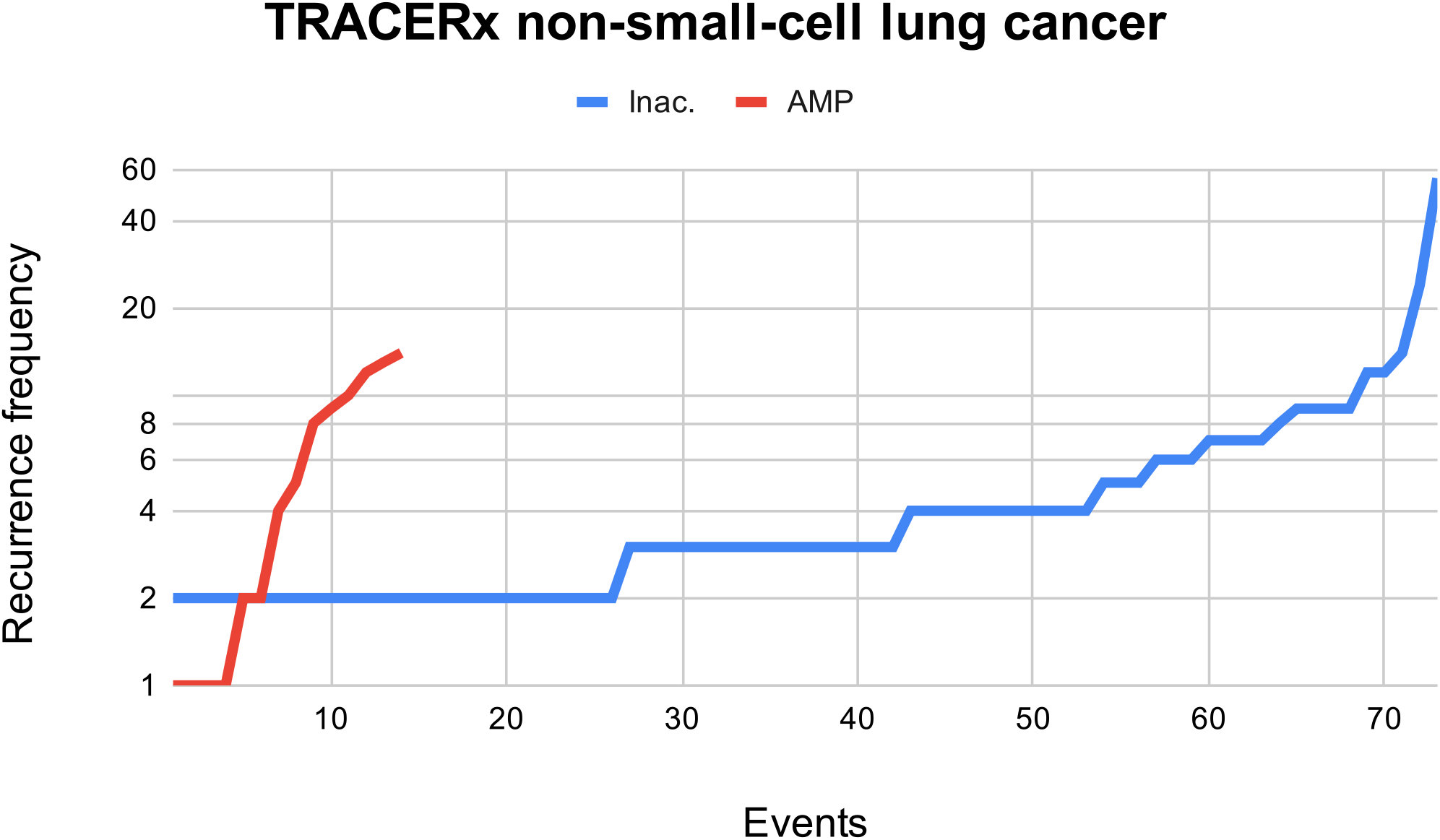
Recurrence frequency of altered genes and chromosome arms in TRACERx lung cohort. The plot shows recurrence frequency of 73 inactivating sequence alteration (blue) and 14 gene amplification (red) events. Events are sorted by their *y* value along the *x* axis for clarity. The *y* axis is in log scale.

**Figure S-4:**
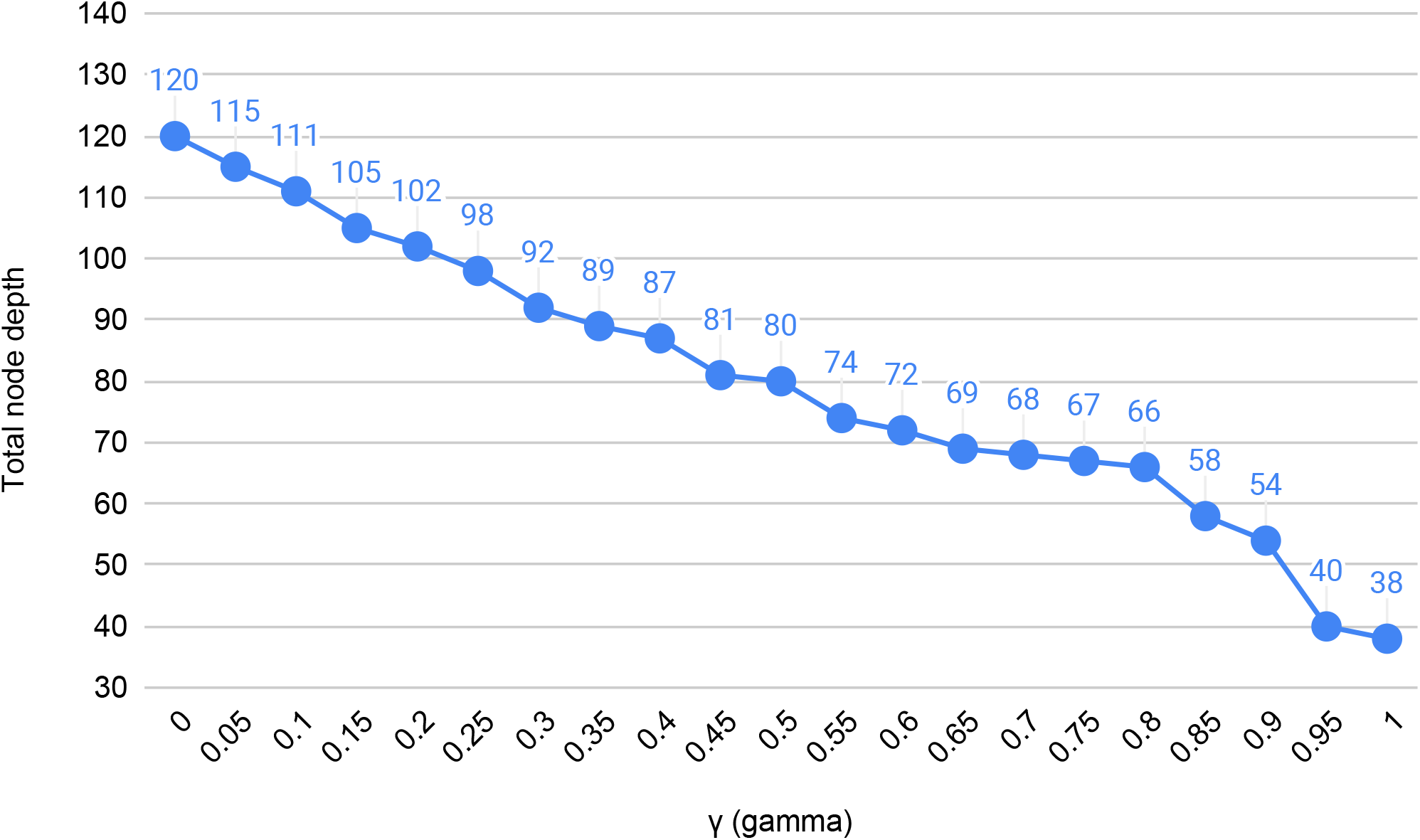
Change of the total node depth in the VHL-rooted conserved evolutionary trajectory tree in ccRCC relative to the change of the *γ* parameter (fraction of tumors that each trajectory needs to be exactly conserved in). Using *t* = 10 (10% of the cohort) and the default value of *δ* = 0.85. The plot shows near-linear negative correlation.

**Figure S-5:**
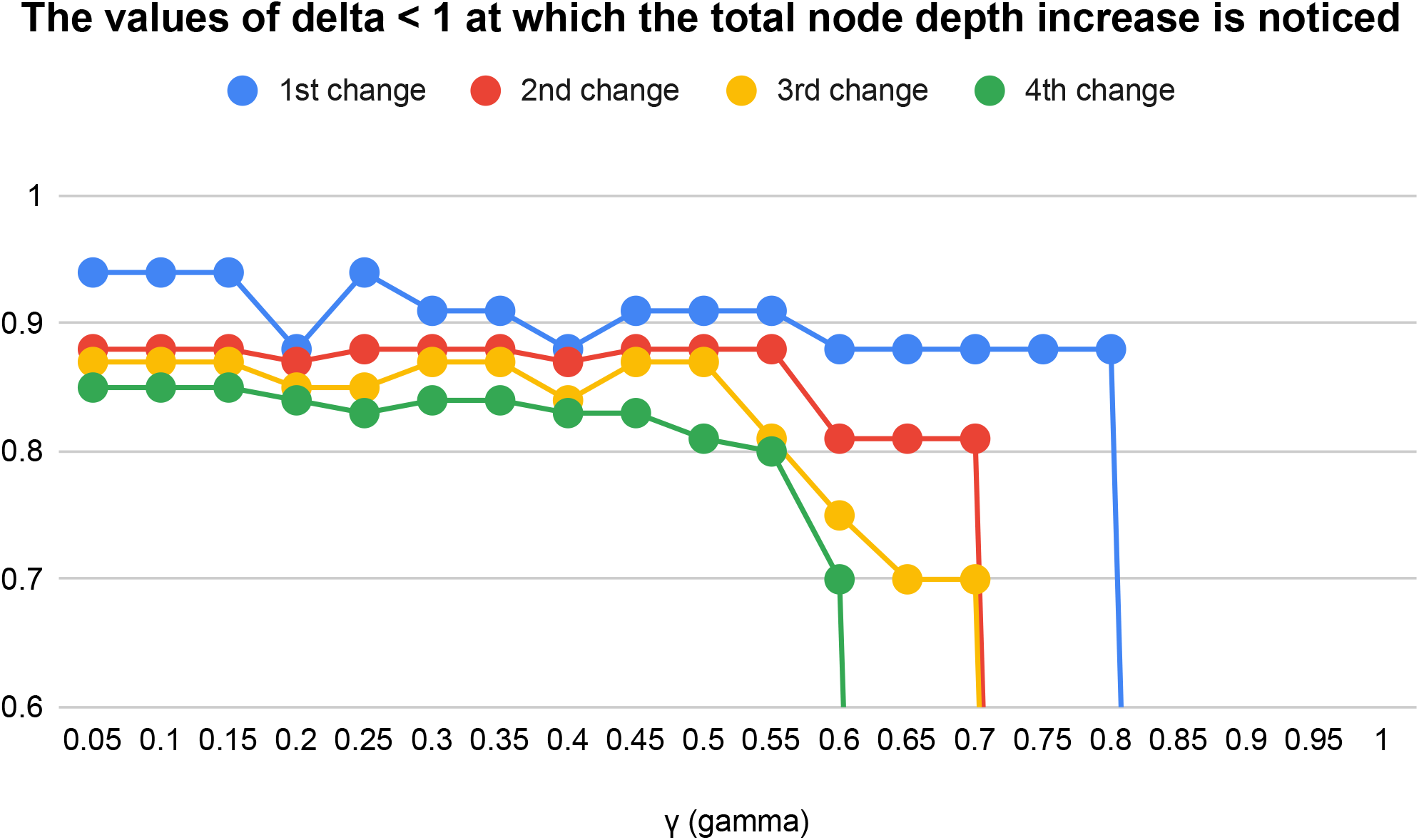
The values of *δ*, smaller than 1, needed to increase the total node depth compared to the previous value of the parameter, in the VHL-rooted conserved evolutionary trajectory tree in ccRCC relative to the change of the *γ* parameter (fraction of tumors that each trajectory needs to be exactly conserved in). Using *t* = 10 (10% of the cohort).

**Figure S-6:**
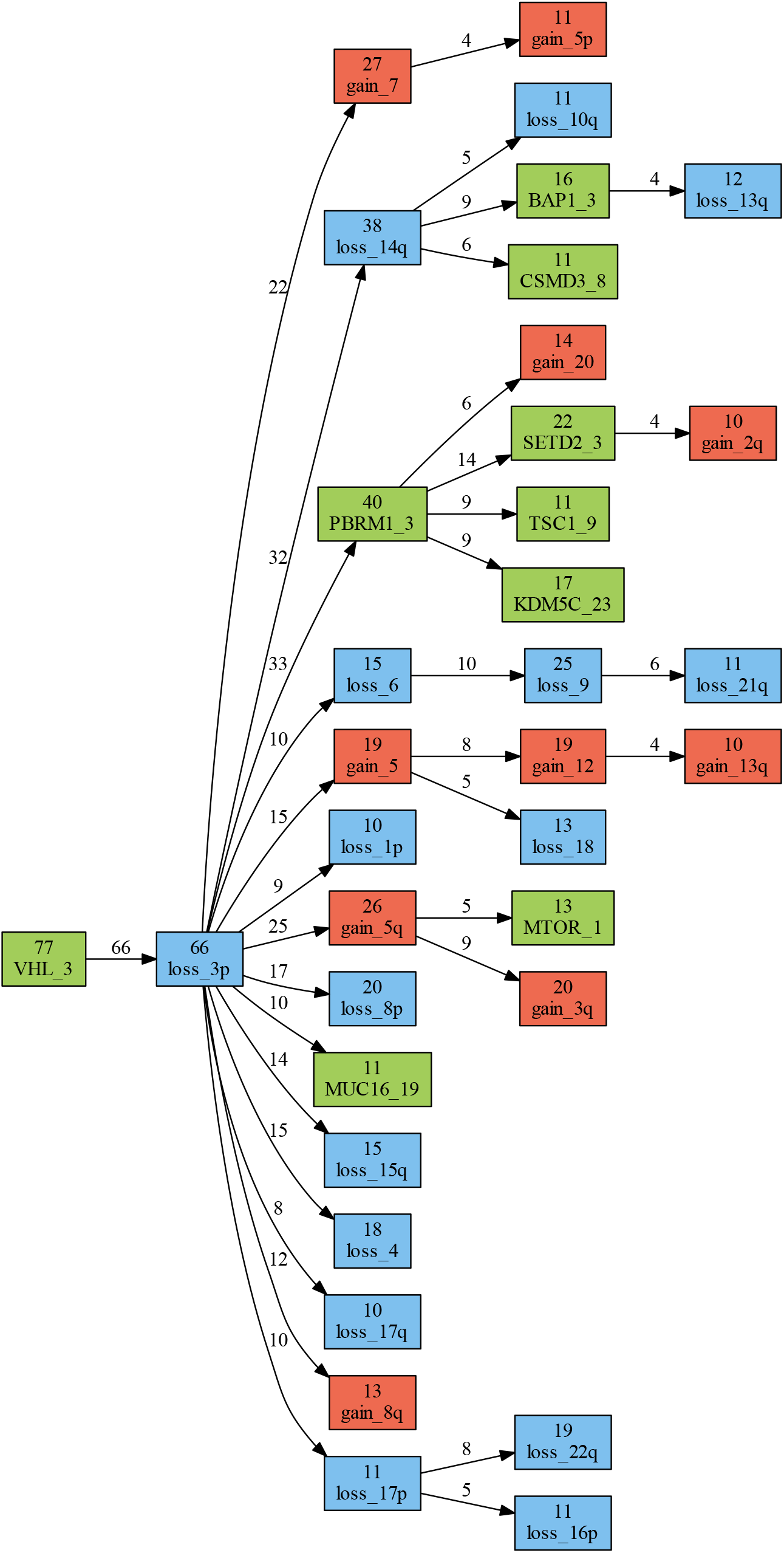
VHL-rooted tree of trajectories with *t* = 10 in TRACERx ccRCC.

**Figure S-7:**
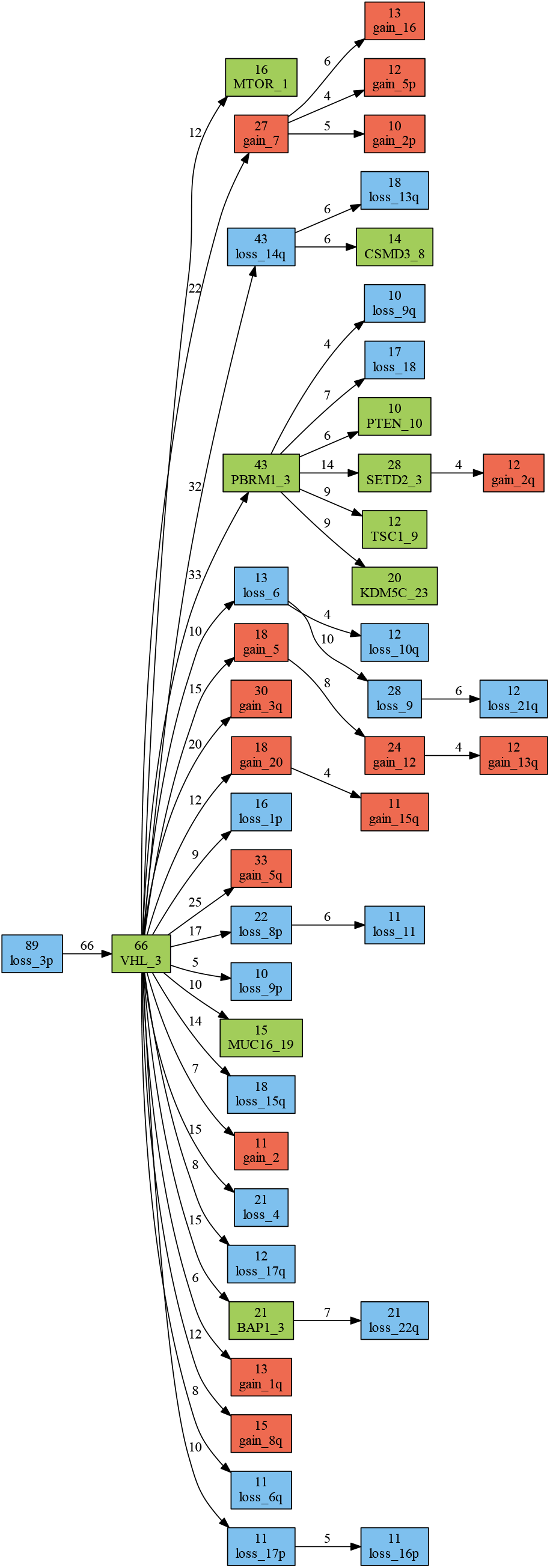
loss3p-rooted tree of trajectories with *t* = 10 in TRACERx ccRCC.

**Figure S-8:**
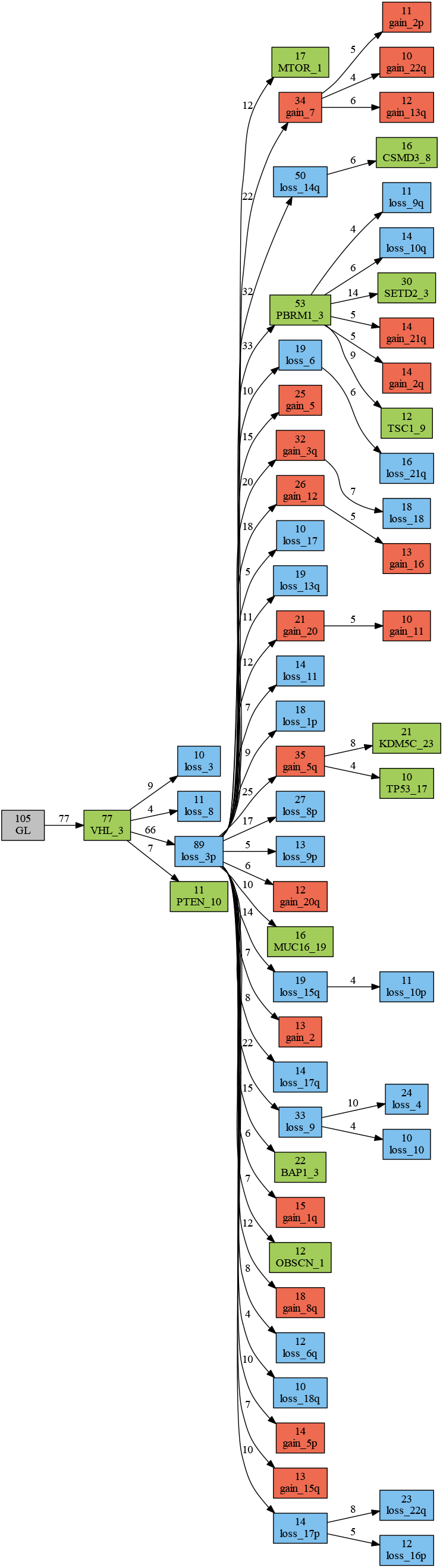
Germline-rooted tree of trajectories with *t* = 10 in TRACERx ccRCC.

**Figure S-9:**
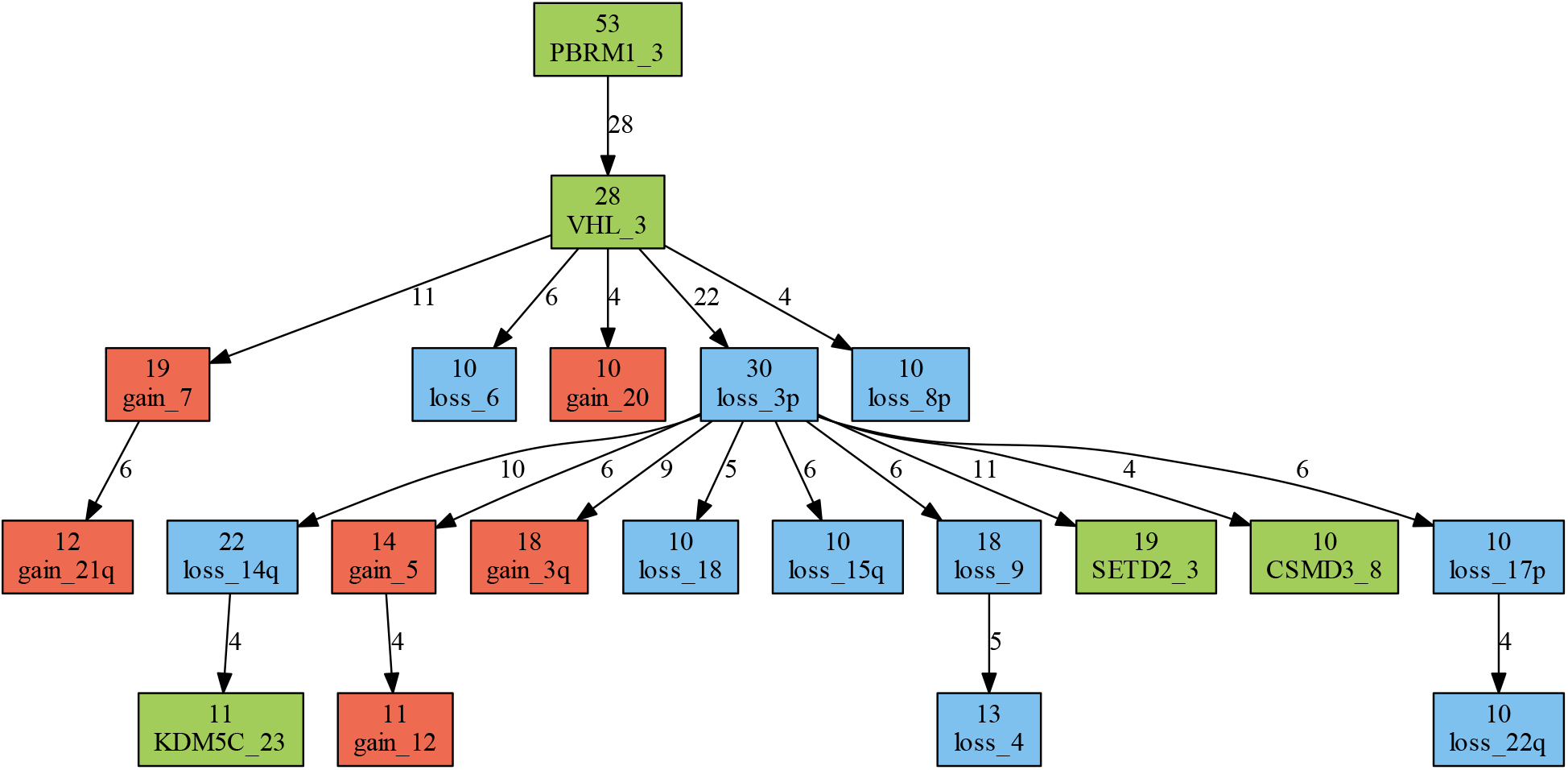
PBRM1-rooted tree of trajectories with *t* = 10 in TRACERx ccRCC.

**Figure S-10:**
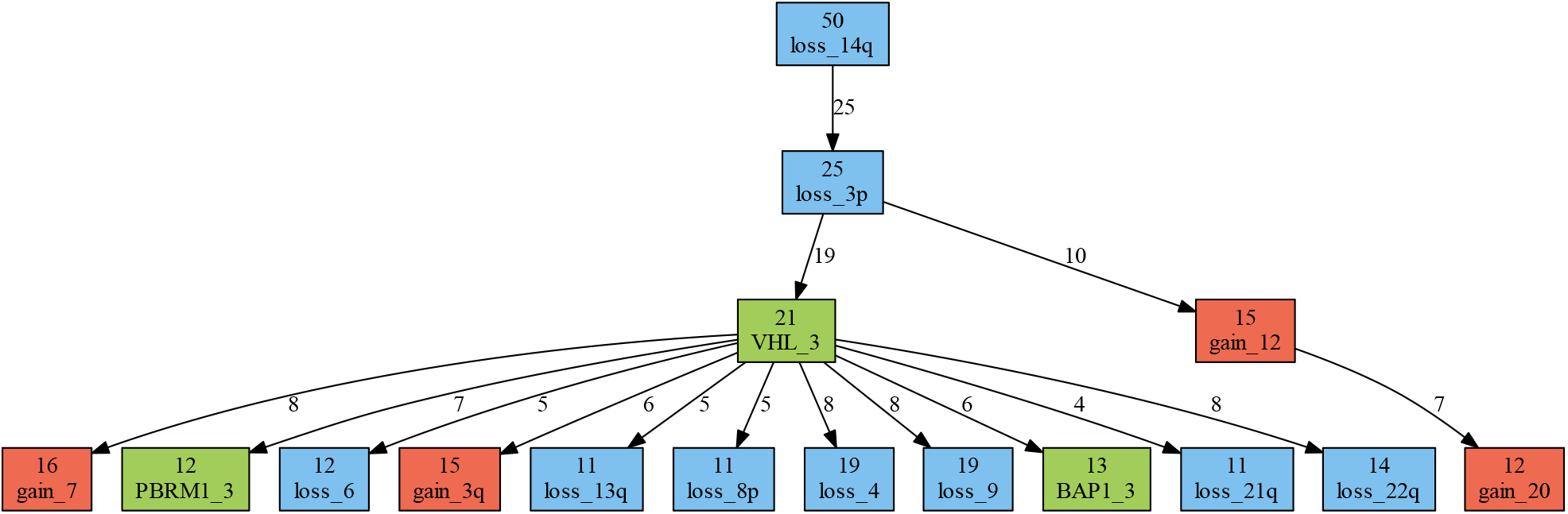
loss14q-rooted tree of trajectories with *t* = 11 in TRACERx ccRCC.

**Figure S-11:**
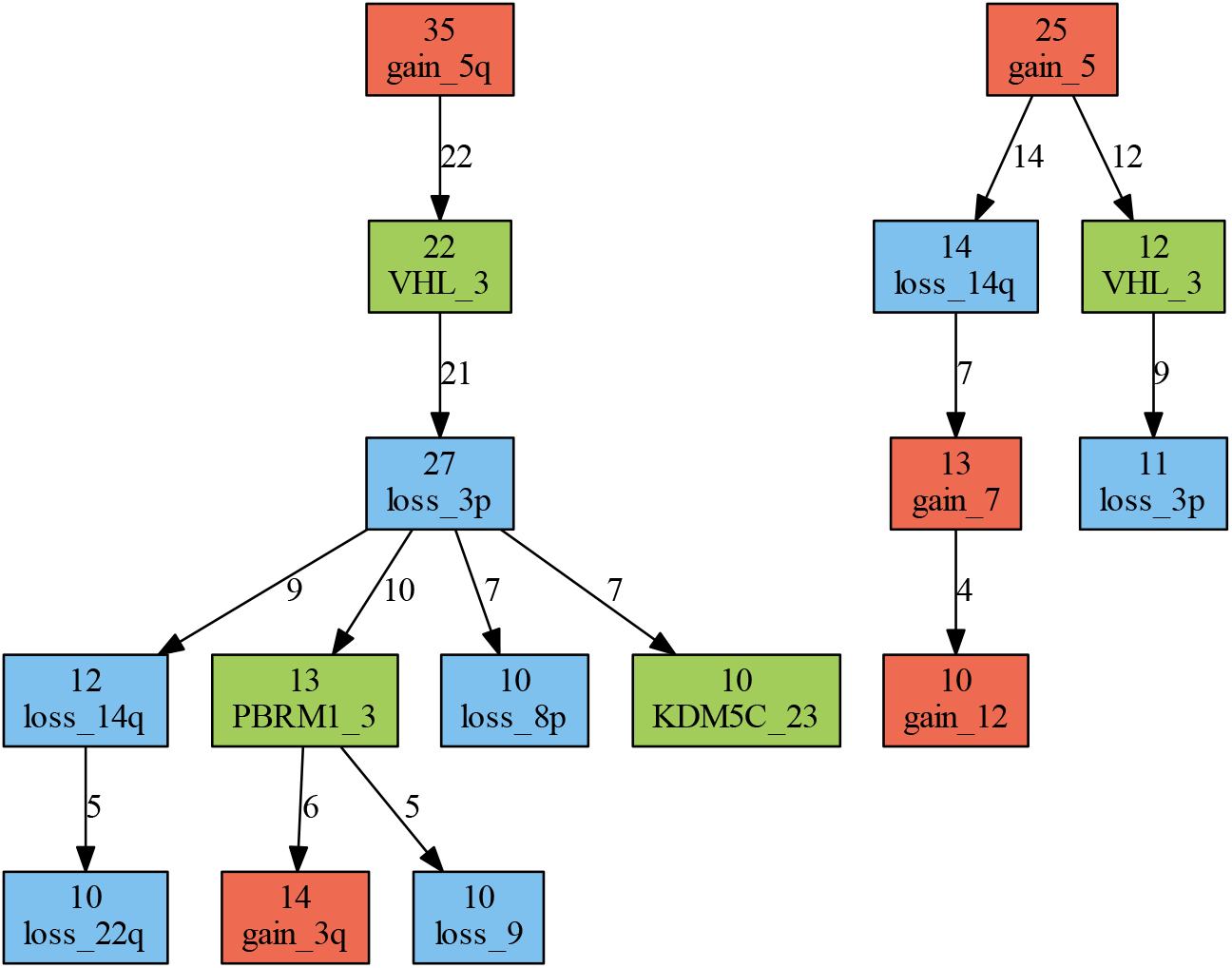
gain5 and gain5q rooted trees of trajectories with *t* = 10 in TRACERx ccRCC.

**Figure S-12:**
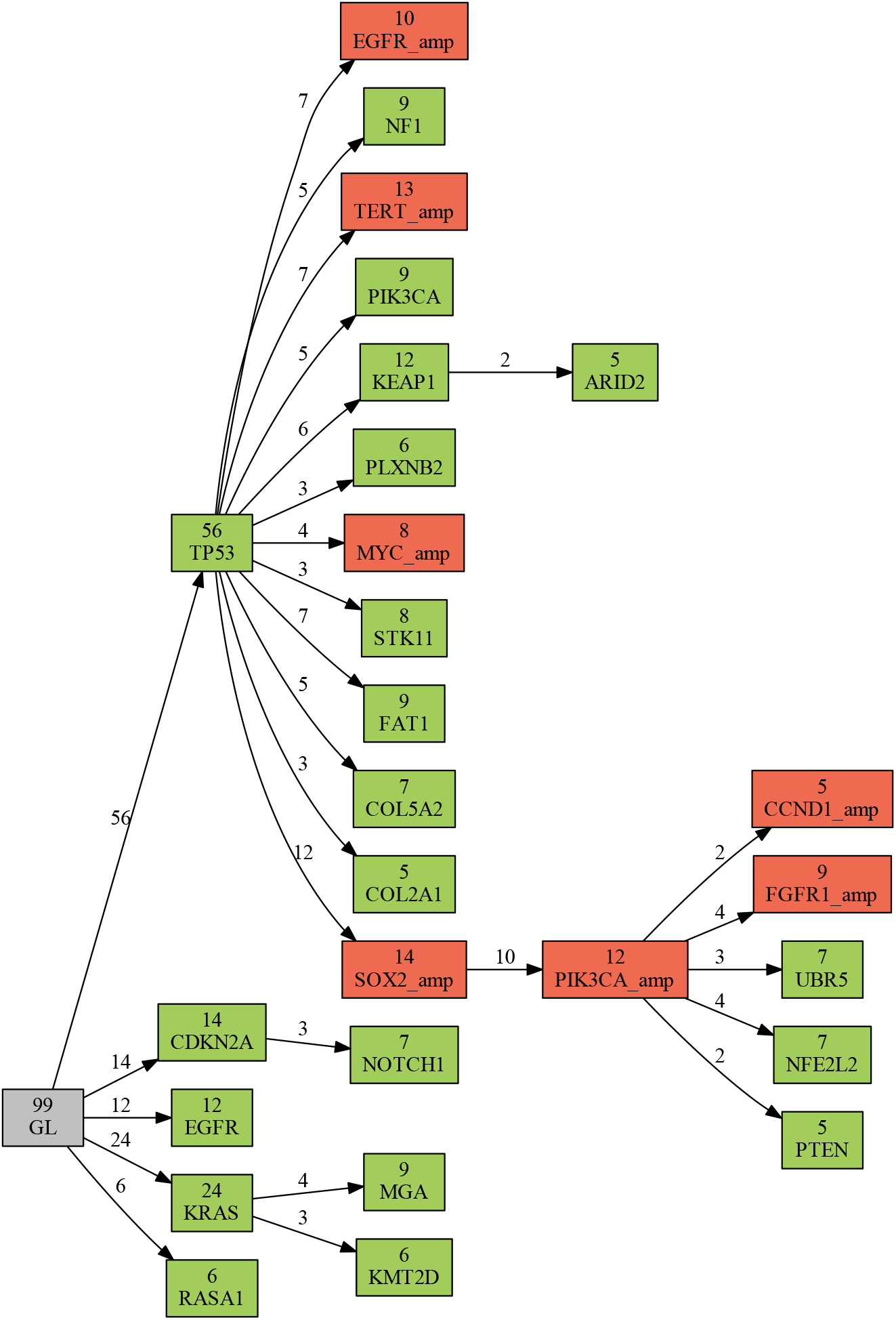
Germline rooted tree of trajectories with *t* = 5 in TRACERx NSCLC.

**Figure S-13:**
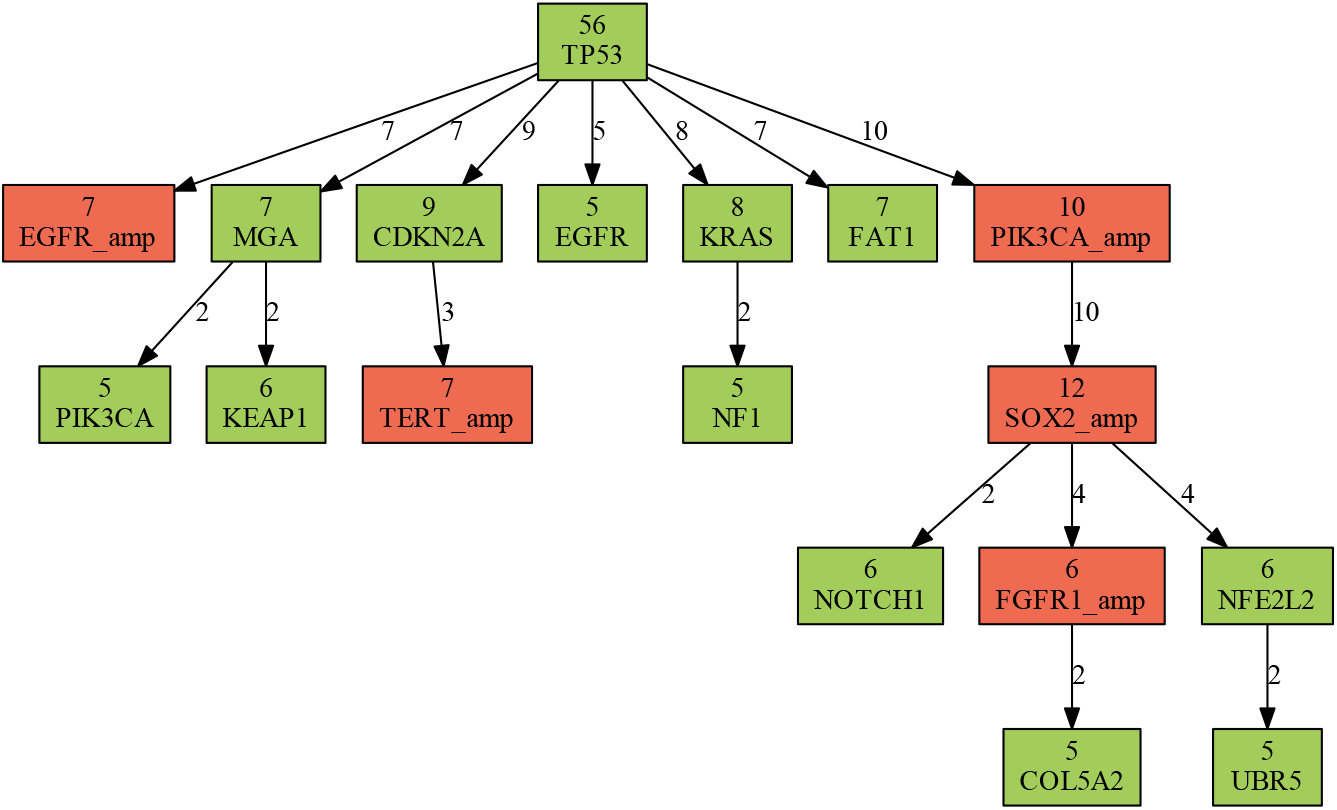
TP53 rooted tree of trajectories with *t* = 5 in TRACERx NSCLC.

## Footnotes

1 We require that the input graph for each tumor should be transitive, i.e. if there are directed edges from *a* to *b* and from *b* to *c* then there must exist a directed edge from *a* to *c*.

2 Note that CONETT allows the user to distinguish distinct types of somatic alterations such as single nucleotide alterations, short insertions, deletions, inversions and duplications within a gene, as well as copy number gains and losses within a chromosomal arm. However, the studies that published the data sets that CONETT was applied to in this paper, did not associate significant biological differences with these distinctions, and thus, for the sake of simplicity and the ability to compare our results, we similarly largely avoided employing such distinctions.

3 This is because the 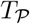 maximizes the total path length.

4 In the case that transitivity of directed edges in the tumor graphs is broken, then *P_u,v_* represents the subset of tumor graphs in which there is a directed path from *u* to *v* (they may lie on a cycle). Then *R_u,v_* represents the subset of tumor graphs in which there is a directed path from *v* to *u* but no directed path from *u* to *v* is present. Since, in general, anti-edges do not need to be explicitly specified, so long as don’t care edges are specified, we may consider *Q_u,v_* to represent the subset of tumor graphs in which there is no directed path between either *u* to *v* or *v* to *u*.

5 As mentioned earlier, CONETT can be set to distinguish alteration types on individual genes or chromosomal arms. We have not treated distinct alteration types differentially in the algorithms described above, however in case certain genes/chromosomal arms included in *S* are not strongly associated with a specific alteration type, (across the tumors) our empirical test can be set to differentiate alteration types as described here. Otherwise *a* can be set to the same value for all nodes.

## Notes

#### Summary of Updates

Substantially re-written Methods Section in order to present our formulation and the method framework more precisely and clearly.

